# Mammalian growth-regulating factors enhance regeneration of recalcitrant transgenic tomato accessions

**DOI:** 10.1101/2025.09.25.678568

**Authors:** Cecile Garchery, Juliette Benejam, Alexia Grau, Justine Gricourt, Esther Pelpoir, Mathilde Causse

## Abstract

Genome editing is now available for many crops. It has increased our ability to study gene function and has changed the field of plant transgenesis. Nevertheless, the ability to regenerate plants from cell culture remains a limiting factor for many crops, and even for species with a good regeneration potential, some accessions remain recalcitrant. The physiological state of plant cells is involved in the process of plant growth and development and is closely linked to the network involving MAP-kinase signaling pathway. Some of the defense genes activated during the cellular repair process of transgenesis show high homologies with mammalian defense genes. We thus compared the percentage of transgenic plants obtained by CRISPR-Cas9 mutation in four genes involved in sugar and acid metabolism after supplementation with different mammalian growth factors and cytokines in six tomato accessions presenting a range of regeneration levels. We demonstrated, through three years of transgenesis experiments, that the use of mammalian growth factors during transgenesis improved regeneration rate of recalcitrant tomato accessions. We demonstrated that using cytokines not only improved transformation of difficult-to-transform accessions but also the production rate of stable secondary lines.

**Summary statement:** Supplementation of transformation medium with mammalian growth-regulating factors enhanced regeneration of tomato recalcitrant genotypes

## Introduction

Since the discovery of the potential of clustered regularly interspaced short palindromic repeat (CRISPR) coupled with the protein Cas9 system (Doudna and Charpentier 2014; Doudna et al. 2018; Gostimskaya 2022), the applications of genome editing has exploded (Rasheed et al. 2021). Genome editing techniques have been shown as essential in various application fields. In plant genetics and breeding research, methods were developed particularly to understand the functional role of candidate genes. Editing techniques make it possible to improve crops more rapidly, without introducing several undesired genes from the donor parent, unlike conventional selection which is long, expensive and can lead to a drop in yield due to linkage drag (Dhugga 2022). The interest of using genome editing for plant breeding has been demonstrated in commercial species (Kausch et al. 2019).

However, in tomato, editing genotypes of interest is often constrained by marked differences in regeneration capacity across genetic backgrounds, highlighting the need to improve this parameter to increase the efficiency of transformation approaches.

Tomato (*Solanum lycopersicum* L.) is a vegetable crop with major agronomical interest. Many breeding programs are conducted to improve tomato yield, yield stability, adaptation to various growth conditions and quality traits (Tanksley 2004; Garchery et al. 2013; Truffault et al. 2018; Schouten et al. 2019; Henri⍰Desaint et al.; Héreil et al.). Improving tomato quality in response to consumer complaints is an important goal of tomato breeding. Sugar and acid contents are among the most important taste-related traits (Causse et al. 2022). The balance between citric and malic acids has been described as critical for fruit quality (Stevens et al. 2007; Bauchet et al. 2017). Using CRISPR Cas9 technology, (Ye et al. 2017) confirmed the functional role of the gene AlMT9, previously identified by GWAS (Sauvage et al. 2014) and its involvement in malate content and fruit quality.

Thus, mastering gene editing in the genetic backgrounds of interest is essential to enable the functional validation of the targeted genes.

The development of plant biotechnology has created great opportunities for tomato plant engineering. Many successes have been recorded in tissue culture and genetic modification of tomato, improving their resistance to environmental stresses and fruit quality and making it a model crop (Gerszberg et al. 2015; Vats et al. 2023b, a). Indeed, tomato has the advantage of proposing many genetic and molecular resources, including several sequenced genomes (Montoya et al. 2002; The Tomato Genome Consortium 2012; Gao et al. 2019; Wang et al. 2019, 2021) and mutagenized populations (Minoia et al. 2010). Moreover, tomato, at least model accessions, can be easily transformed by several tools (Agrobacterium-mediated, direct DNA uptake via protoplasts, biolistic, *in planta* Particle Bombardment (Imai et al. 2020; Vats et al. 2023b).

The CRISPR Cas9 technology has been developed rapidly on tomato since 2014 (Brooks et al. 2014; Lor et al. 2014; Rothan et al. 2019). Tiwari et al (2023) presents the long list of genes studied by CRISPR technology on tomato. In Japan, where the marketing of modified organisms is authorized, the first tomato edited by Crispr/Cas9 to increase its GABA content was commercialized in 2022 (Waltz 2021). Tomato transformation is generally carried out on a limited number of accessions (Maren et al. 2022). It has been reported that the frequency of regeneration in tomato transformation varies from 6% for ‘Pusa Ruby’ (Sree Vidya et al. 2000) to 40% for ‘Micro-Tom’ (Sun et al. 2006; Qiu et al. 2007).

Regeneration frequency in tomato genetic transformation refers to the percentage of in vitro cultured explants that develop regenerated shoots (full or developing plants) after the introduction of a transgene or genome editing system (such as CRISPR-Cas9).

One of the main limitations affecting most transformation methods is the ability of plants to regenerate (Ikeuchi et al. 2016; Kareem et al. 2016; Méndez-Hernández et al. 2019). The ability of plant regeneration by *de novo* organogenesis or somatic embryogenesis is fundamental but far from being elucidated. It is therefore becoming necessary to understand the factors limiting the regeneration of certain genotypes in order to extend this method to a large pool of genetic diversity, and more particularly to varieties with commercial interest, as genome editing techniques become essential tools for plant breeding.

The ability of genotypes to regenerate depends in part on the developmental plasticity and totipotency of plant cells. Several genetic and epigenetic mechanisms involved in the ability or induction of cell regeneration have been identified (Iwase et al. 2015; Fan et al. 2015; Méndez-Hernández et al. 2019; Debernardi et al. 2020) but the molecular mechanisms involved are not yet well understood. Perez-Garcia and Moreno-Risueno (2018) show that there are different stem cell types and a large number of regulators linked to the fate of these cells. Other recent studies suggest that species-specific environmental stimuli (biotic and abiotic) may be required for cell reprogramming in relation to known cell repair mechanisms (Sugimoto et al. 2019; Ikeuchi et al. 2019; Ichihashi et al. 2020). These results suggest that there are genotype-specific mechanisms associated with cell repair or stem cell differentiation which could explain differences in the transformation capacities of certain genotypes (Long et al. 2022). Overexpression of some genes encoding kinases during embryogenesis has been shown to increase the success rate of regeneration (Banno et al. 2001; Debernardi et al. 2020). In mammals, there are three large families of transmembrane receptor kinases involved in cellular regeneration mechanisms: tyrosine kinase receptors (TKR), serine threonine kinase receptors (STKR) and histidine kinase receptors (RHK) (Rehli 2002; Carpenter and Liao 2009; Combarnous 2013). In both mammals and plants, cell growth regulators are central molecules in cell signaling (Chevalier and Walker 2005; Clos 2012) and can initiate regulatory cascades, including the activation of protein kinases and transcription factors.

The integration of environmental signals at the cellular and tissue level is essential in plants, a mechanism in which CLE peptides, notably CLAVATA3, and their CLAVATA-like receptors play a role that has now been demonstrated (Bashyal et al. 2024). Overexpression of the growth factor gene GRF5 of Arabidopsis has been shown to promote transformation in both dicotyledonous and monocotyledonous species (Kong et al. 2020). The regulation of kinases by growth regulators (Lu et al. 2020; Bull et al. 2023) could therefore be an interesting lever for improving plant regeneration capacity and a tool that can be used to compensate for blocking environments.

A number of plant defense systems analogous to animal systems have been described for a long time in plants (Bergey et al. 1996; Ryan and Pearce 1998; Ryan 2000). A major advance in these studies is the discovery of the wound signaling system, which orchestrates the defense responses triggered by these damages in an analogous manner to cytokines in animal immunity. These secondary endogenous danger signals in plants, which modulate the immune response, should therefore be called phytocytokines (Gust et al. 2017; Wang et al, 2025).

In this study, we hypothesized that tomato recalcitrant accessions are deficient or limited in growth factors capable of activating kinase-type regulatory pathways. Among the three major families of kinases involved in cell regeneration mechanisms, no plant-derived growth regulators are currently available on the market but only those of murine or human origin. We thus tested if an exogenous supply of mammalian growth regulators could improve the transformation capacity of recalcitrant tomato varieties. The ability of three growth regulators to improve transgenesis was compared on the transformation success of six varieties for the modification of four genes linked to sugar and acid metabolism.

## Materials and methods

### Plant materials

Transgenesis experiments were carried out on six tomato accessions: *‘Cervil’*, ‘Levovil’, ‘LA1420’, ‘Criollo’, ‘Ferum’ and ‘Stupike’. The six accessions were selected to represent genetic and phenotypic diversity for fruit quality (Pascual et al. 2013). The accessions *Criollo* and *LA1420* have been described as having a low malate content in the fruit, while Levovil has a higher content (Bauchet et al. 2017). *Cervil* has a high sugar content in the fruit, on the contrary to Levovil (Albert et al. 2018). The accessions Levovil, *LA1420* and Criollo are considered easy to transform and and Stupike, *Cervil* and Ferum as difficult to transform or even recalcitrant to transgenesis for the *Cervil* accession.

### Gene design of Crispr Cas9 constructs

Transgenesis experiments were done over three years (2018, 2019 and 2022) using four genes as mutation targets: the Mal30 gene (Solyc06g072930) coding for a function still unknown but probably involved in malate transport, an aluminum-activated malate transporter (Bauchet et al. 2017) ; the FRK3 gene (Solyc02g091490) encoding a fructokinase 3 for which two constructs have been made and used (Knock out and Knock in), and considered as candidate gene for sugar content (Albert et al. 2018); the Invertase 3 (suc3) gene (Solyc03g083910) encoding a vacuolar invertase localized on chromosome 3, also modulating sugar content (Chetelat et al. 1995; Albert et al. 2018) and the Invertase 9 (Lin 5) gene (Solyc09g010080) encoding an apoplastic invertase located on chromosome 9 (Zanor et al. 2009) (Supplementary Table 1a, b).

*Agrobacterium tumefaciens* strain ‘C58’ and the helper plasmid ‘pGV2260’ were used as described in Hamza & Chupeau (Hamza and Chupeau 1993). Two cloned plasmids were redesigned for CRISPR/Cas9 technology. A ‘pDE-Cas9 nickase’ plasmid was used to produce knock-out (KO) mutants. The second plasmid “pDicAID-Cas9 nickase” fused to a cytidine deaminase was used to produce base editing mutants of the FRK3 gene from experiment 2022. To target the sequence of interest, two guide RNAs (gRNAs) for KO and one for base editing were designed for the targeted genes and integrated into the respective plasmids. The complete plasmid sequences, including the sequences of the selection genes encoding the antibiotics, are available in **Supplementary Table 2**.

### Selection of growth regulators by sequence homology analysis

To determine mammalian growth regulators that could be tested on tomato accessions, we studied the homology of the protein sequences of four receptors representing three mammalian receptor families with tomato protein sequences, using the computer program UniProtKB (Universal Protein Resource and Functional Database). Accordingly, the study focused on one human receptor, Transforming Growth Factor Receptor (TGFR) and three murine receptors, the Murine Epidermal Growth Factor Receptor (EGFR) and two murine interleukin 1 and 2 receptors (IL1R1 and IL1R2). We therefore used the regulatory ligands, Transforming Growth Factor-β1 (TGF-β1) which activates the STK receptor pathway, Epidermal Growth Factor (EGF) which activates the RTK receptor pathway and interleukin 1β (IL1-1β) which activates the RHK receptor pathway. Blasts (Basic Local Alignment Search Tool for Proteins) using mouse EGFR (query: Q01279), human TGF R1 (query: P36897) and murine IL-1R1, IL-1R2 (query: P13504 and P27931) as queries against *Solanum lycopersicum* species-specific protein sequence databases allowed us to identify homologies. The protein sequences of the ligands are available in Supplementary Table 3.

### Plant transformation and growth conditions

A single colony of *Agrobacterium tumefaciens* was inoculated in LBA medium (Bertani Giuseppe 1951) and incubated with antibiotics rifampicin 50 mg/L, ampicilin 50 mg/L, spectinomycine 100 mg/L for 48 h at 28°C. Colony growth was followed by seeding until an OD_600 value equal to 0.4 was reached. The bacterial culture was centrifuged and suspended in culture medium supplemented with thiamine, 2-4 D and acetosyringone (co-culture). Sterile cotyledon sections from about a hundred plants per accession were placed on the solid coculture medium containing 30 µL of bacterial suspension supplemented with 20 µL of regulatory solution (or water for control) containing the cytokine growth factors. We used transforming growth factor β1 (human recombinant TFG-β1 factor), epidermal growth factor (murine recombinant EGF factor) and a murine paracrine pro-inflammatory factor, interleukin (IL-1β), each at a final concentration of 2 µM. The concentrations for use are recommended by the company PEPROTECH (Paris – France) from which we purchased the three growth factors.

For each plasmid construct, 6 to 10 Petri dishes, each containing an average of 10 explants, were generated. Explants were incubated in the dark for 48 h, then transferred to a selective MS medium (Murashige and Skoog 1962) for 10 weeks in a climatic chamber (16h/8h light, 20/16°C and 50% humidity). For the first few days, explants were protected by a UVB filter prior to penetration and following the application of radiation-sensitive growth factors (300 nm). The support medium was renewed every 14 days. Regenerated shoots were isolated and transferred to a nutrient-depleted root induction medium composed of 1/2 MS (Hamza and Chupeau 1993). The composition of all culture medium is detailed in Supplementary Table 4.

The regenerated primary seedlings were analyzed and selected by sequencing (Sanger technique) for the mutated part of the gene of interest. The selected mutant plants were acclimated and grown in a greenhouse to produce seeds by self-fertilization. The secondary homozygous mutants were used for the qPCR experiments.

### Scoring regeneration and evaluation of regulator performance

Regeneration performance was estimated by assessing cell/explant quality at five growth stages: (1) cell clustering and callus formation, (2) bud differentiation, (3) isolated plantlets, (4) acclimated plantlets, and (5) edited lines obtained. Scores were collected on 10 plates/explant by three different people qualified in cell biology. The scoring scales are detailed in Table 1. For the last step, the presence of the mutation in each mutant line was validated by sequencing. Leaves were sampled on each isolated plantlet or plants acclimated in the greenhouse and DNA was extracted using the DNeasy 96 Plant kit (QIAGEN). A PCR was performed using primers specific for the corresponding target gene, and the amplicons were sent to GenoScreen (France) for sequencing (Genome editing results of the secondaries of the *Cervil* and Levovil accessions obtained with the addition of TGF during transgenesis for the target genes Suc3, FRK3 and Lin5 are listed in Supplementary Table 5).

**Table 1.**
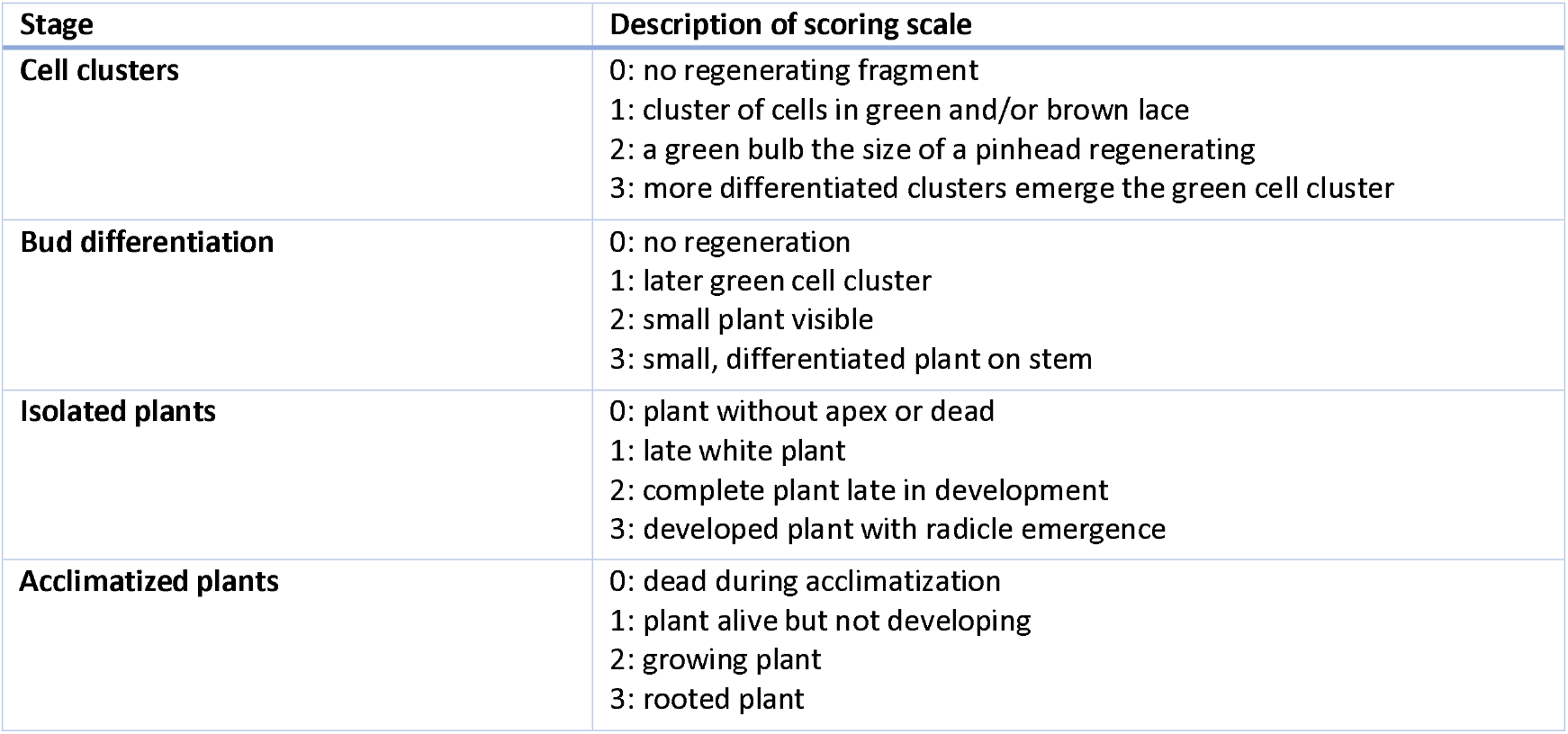
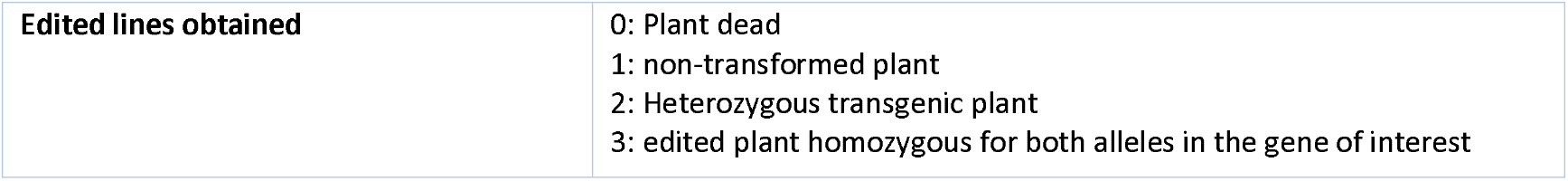
Scoring scales used at each regeneration stage.

Regeneration performance was estimated by assessing the quantity and quality of regenerated plants. Quantity was transformed to frequency of in vitro-cultured explants that developed regenerated shoots (complete or developing plants) after the introduction of a genome-edited transgene (CRISPR-Cas9 in our experiments). We noted regeneration frequency (%) = (Number of explants that produced shoots / Total number of inoculated explants) × 100.

Furthermore, a gene expression analysis was carried out on secondary plants to verify whether the transgene, integrated thanks to the growth factors used during transgenesis, was maintained in a stable manner and transmitted to subsequent generations. The expression of three genes (*Suc3, Lin5 and FRK3*), was assessed by RT-qPCR in mutant lines of the *Cervil* and Levovil genotypes generated during the 2019 experiment. RNA was extracted from the same samples as DNA extraction using the Spectrum Plant Total RNA kit (Sigma-Aldrich). cDNA was produced by retrotranscription after RNA normalization to a concentration of 5 μg/μL using fluorometry for assay (Invitrogen Qubit 3.0 fluorometer). RT-qPCR was performed on three replicates per line and a negative control (water) using Brilliant II SYBR® Green QPCR Master Mix enzyme (Agilent). To compare expression results, a pool of all cDNAs was added to the experiments, estimating variability between RT-qPCR cycles.

Details of the constructs, accessions, growth factors and phenotyping traits evaluated in each experiment are summarized in **Table 2**.

**Table 2.** Details of constructs, accession and phenotyping traits evaluated in each experiment.

| Year | Type | Accession | Target Gene | Regulator | Phenotyped trait |
| --- | --- | --- | --- | --- | --- |
| 2018 | KO | Cervil, Levovil, Ferum | Malate 30 (SolyC06g072930) | TGF-beta1 vs control (sham inoculated) | Regeneration yield |
| 2019 | KO | Cervil, Levovil, Stupike, Criollo, LA1420 | Malate 30 (SolyC06g072930) | TGF-beta1 vs control (sham inoculated) | Regeneration yield |
| 2019 | KO | Cervil, Levovil | Invertase 3 (solyC03g083910)<br>Invertase 9 (Lin5 SolyC09g010080)<br>Fructokinase 3 (FRK3 SolyC02g091490) | TGF-beta1 vs control (sham inoculated) | Regeneration yield<br>Gene expression of the three genes |
| 2022 | KI | Cervil | Fructokinase 3 (FRK3 SolyC02g091490) | TGF-beta1, EGF, interleukine1beta vs control (sham inoculated) | Regeneration yield |

### Statistical analysis

Statistical analyses were carried out to test the differences observed between the control and treated lines. After verification of the normal distribution (using Shapiro’s test) and homogeneity of variances (using Bartlett’s test), we compared means (Student’s parametric test) of mutant construct versus control construct, when the normal distribution of the data allowed it. Otherwise, non-parametric Wilcoxon-Mann-Whitney tests were used to make two-to-one comparisons (presence vs absence of regulators).

In order to analyze the qPCR results, the following calculations were performed:

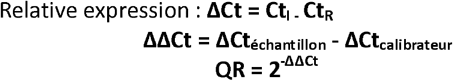

With Ct the Cycle Threshold (cycle during which the PCR curve crosses the threshold), I the gene of interest, R the reference gene and RQ the relative quantification. For these analyses, the calibrator corresponds to the test range point diluted to 1/5 (same dilution as the samples). All these calculations were performed on the average Ct of the 3 technical replicates. The RQ was then expressed in log2 for ease of reading.

Tukey tests were also performed when a comparison of all mutants was deemed relevant. Analyses were carried out genotype by genotype, followed by an overall comparison of results. All these data were then compiled in box-plot, showing standard deviations and significant differences. All statistical tests were performed using RStudio software, and the error threshold was set to α = 0.05.

## Results

We studied the impact of supplementation with three ligands specific to the three transmembrane domain receptor families of the tyrosine, serine/threonine and histidine kinase signaling pathways. We used (i) two long haul cytokines TGF-β1 “Transforming Growth Factor-β1 recombinant human”, (ii) EGF “Epidermal Growth Factor Recombinant Murin” activating single-pass transmembrane receptors of the catalytic domains of the serine/threonine and tyrosine pathways and (iii) a paracrine pro-inflammatory factor, recombinant Murine interleukin IL-1β, specific for receptors directly coupled to kinases or other proteins and lacking a catalytic domain, activating the histidine kinase pathway (sequences in Supplementary Table 3).

### Sequence comparison between mammalian and tomato growth factor proteins

In order to identify growth factors of human or murine origin that could have an effect in tomato, we first looked for protein sequences in tomato that were homologous to mammalian regulatory factor receptors of the different growth factors.

#### TGF-β1 receptor

The results of pBlast performed on the human receptor sequence (Supplementary Figure S1a), P36897 TGFR1_HUMAN (503 amino acids) as a query against *Solanum lycopersicum* protein sequence databases) found 250 potential homologies. The most significant homology corresponded to the protein kinase domains in tomato.

Among the top 20 ranked potential proteins according to UniProt, a kinase domain-containing protein (Solyc03g114310) showed high similarity to a human receptor sequence (Supplementary Figure S1b), with a score of 108 bits (E-value 4.10^-26^). It had 29% identity and 48% similarity, indicating notable amino acid conservation, despite gaps of 19%, suggesting structural divergence (Supplementary Figure S1c).

Two other interesting proteins suggested by Uniprot (Solyc02g072070 and Solyc04g051510), also showed significant similarities (Supplementary Figure S1d, S1e). These proteins are involved in the activation of the tomato serine/threonine kinase receptor, itself involved in the regulation of genes linked to plant development in response to brassinosteroids and in the perception of hormonal signals linked to the defense of the injured plant (Montoya et al. 2002; Nie et al. 2019; Wang et al. 2019, 2021).

#### EGF receptor

The results of pBlast carried out using the murine EGF receptor sequence identified 250 potential homologs in *Solanum lycopersicum* (Supplementary Figure S2a and S2b). The first identified protein contained a kinase domain (Solyc10g055720) and had high similarity (with a score of 148 bits and an E-value of 3.10^-37^) with 34% identity and 55% positive similarity (Supplementary Figure S2c). Furthermore, among the proteins with homology, the protein kinase Q41328 (Solyc12g098990) Pto interacting protein 1 was characterized by (Zhou et al. 1995) as a functional protein of the tomato catalytic site involved in tomato immune responses (Supplementary Figure S2d).

#### Interleukins Receptors

We found two specific receptors for the growth regulator interleukin (IL-1β) whose protein sequences are shown in Supplementary Figure S3a (IL1R1_MOUSE Interleukin 1 Receptor 1 P13504) and in Supplementary Figure S4a (IL1R2_MOUSE Interleukin 1 Receptor 2 P27931), according to UniProt databases.

Among the 12 pBlast results from the IL1R1_Mouse receptor interleukin 1 receptor P13504 query (Supplementary Figure S3b supp) against the *Solanum lycopersicum* protein sequence database, we noted one protein (Solyc08g083300) involved in the recognition and hydrolysis of the peptide bond at the C-terminal Gly of ubiquitin and also involved in the processing of polyubiquitin precursors as well as ubiquitinated proteins (The Tomato Genome Consortium 2012). There was a significant similarity with a moderate score of 33.1 bits and an E-value of 0.54. The positive identities and similarities were 32% and 46%, respectively, and showed a moderate level of amino acid conservation between the sequences (Supplementary Figure S3c).

Among the 8 pBlast query results with IL1R2_Mouse Interleukin 1 receptor 2 P27931 compared to the *Solanum lycopersicum* protein sequence database (Supplementary Figure S4b), one uncharacterized protein (Solyc09g010480) showed a moderate level of similarity (34.7 bits and E-value of 0.029). The 23% identity and 40% positive similarity indicated that although some regions of the proteins are conserved, there are also notable differences. These results could indicate that the compared proteins share similar functional motifs or domains, but they may also have distinct functions (Supplementary Figure S4c).

### Effect of growth factors on different tomato accessions: regeneration yield

#### Diversity of regeneration between accessions

Figure 1a presents the diversity of regeneration level of six accessions, at each step of regeneration, without addition of any growth factor. The accessions *Cervil, Ferum, Stupike* and *Levovil* presented extremely low or even zero levels of regeneration, while *Criollo* and *LA1420* were among the most efficient accessions.

**Figure 1:**
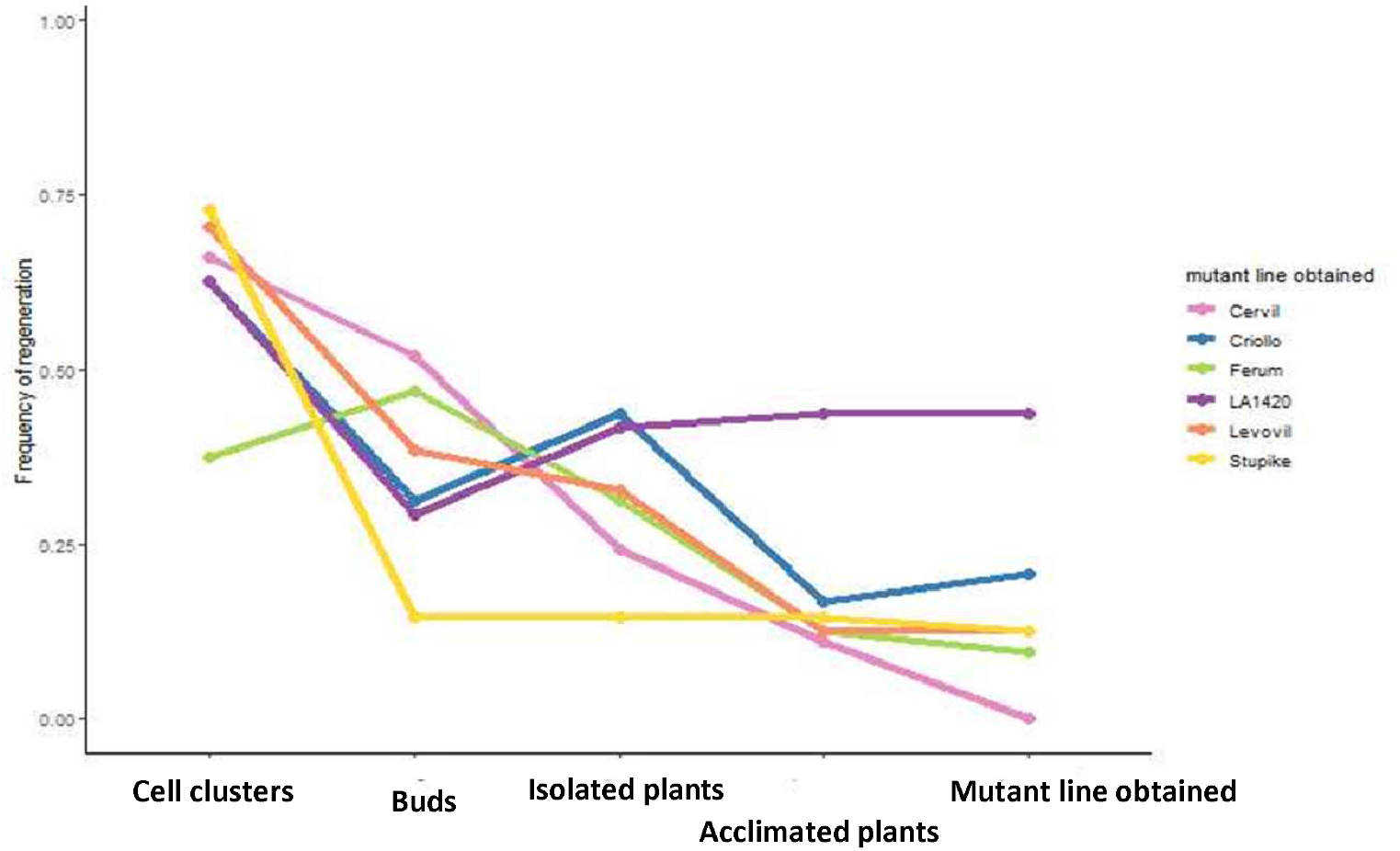
Comparison of the regeneration levels of six tomato accessions during the transgenesis process without supplementation of growth regulators at five stages of regeneration (cell clusters, buds, isolated seedlings, acclimated plants and mutant plants obtained). Each point corresponds to the average over 3 years of experiment.

The *Cervil* accession was considered as recalcitrant to transgenesis with very few acclimated plants not any edited despite a satisfactory capacity to return to the undifferentiated cell stage. The *Ferum* accession had a low capacity to return to an undifferentiated embryonic state. *Cervil* and *Ferum* showed a correct re-differentiation rate compared to all other accessions at the same stage but lost this advantage at later stages of differentiation, tissue organization and organ formation. Levovil and *Stupike* showed weaknesses at the re-differentiation stage (*Stupike* essentially) and showed difficulty in organizing tissues and forming differentiated organs. The “high-performance” accessions, *Criollo* and *LA1420*, also encountered difficulties at the re-differentiation stage, but managed to organize functional tissues. However, at the stage of organ formation (mainly root system) and acclimatization, *Criollo* showed its regeneration rate decreasing. *LA1420* was the optimal candidate for regeneration in the context of transgenesis with efficient integration of the transgene.

To test the hypothesis that cellular signaling and communication systems, similar to those in plants and mammals, provide advantages during the transgenesis process, we conducted a series of experiments. During the first year, we tested the effect of addition to the medium of a growth regulator (TGF-β1) on two accessions (*Cervil* and Levovil). During the second year, we tested the effect of TGF-β1 on six accessions (*Cervil, Levovil, Criollo, Ferum, Stupike, LA1420*) and during the third year, we evaluated the effect of three growth regulators (TGF-β1, EGF, and IL-1β) on the recalcitrant accession to transgenesis in *Cervil*.

To summarize our results, we will present successively: (1) the quantitative and qualitative effect of the growth regulators on *Cervil*, the most recalcitrant accession, to transgenesis; (2) the qualitative response to the TGF-β1 regulator of six accessions and (3) the beneficial effect of the three growth regulators (TGF-β1, EGF and IL-1β) on the recalcitrant accession *Cervil*.

#### Effect of TGF-β1 on the recalcitrant Cervil accession

Based on data collected over three consecutive years, supplementation of the growth factor TGF-β1 resulted in a significant increase in the regeneration frequency of transformed calli of the *Cervil* accession compared to the control group that did not receive TGF-β1 (Figure 2a). The effect of TGF-β1 is particularly strong at the cell cluster stage, which suggests that its action begins with the dedifferentiation stage towards a more embryonic or undifferentiated state of the explants.

**Figure 2:**
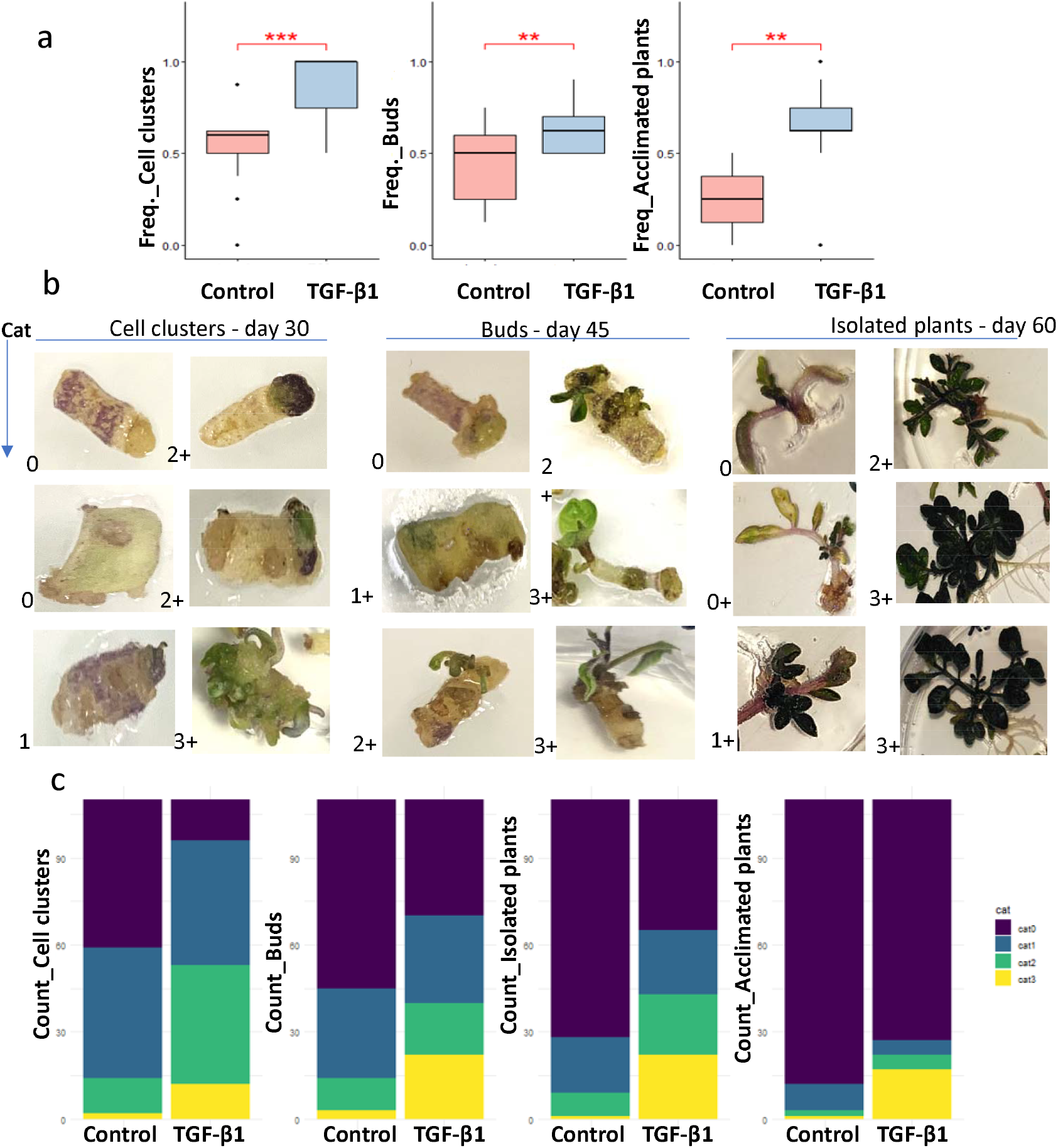
Effect of TGF-β1 on the regeneration of the recalcitrant *Cervil* accession (over three years of experiments : 2018 Mal30-KO gene mutation – 2019 Mal30-KO, Suc3-KO, FRK3-KO, Lin5-KO gene mutation – 2022 FRK3-KI gene mutation) for three stages of regeneration (cell clusters, buds and isolated plantlets) with the supplementation of TGF-β1 at a concentration of 2 mg/mL compared to the control condition (control) without addition of growth factor. **a :** Comparison of the frequency of regeneration of transformed calluses at each stage. ^**^ p<0.01; ^***^ p<0.001. **b :** Visual representation of the quality scores of explants transformed into regeneration at three stages throughout the 60 days of development process. Four quality categories were scored : 0 - absence of regeneration; 1 - little visible characteristic of regeneration; 2 - visible characteristic of development; 3 - complete and optimal regeneration. + indicates whether the explant is supplemented with regulator TGF-β1. **c :** Comparison of the number of regenerated calli in *Cervil* at four developmental stages (cell clusters, buds, isolated seedlings, and acclimated plants), with or without TGF-β1 supplementation (2 mg/mL). Callus quality is divided into four categories: Cat0 (purple) – no regeneration, Cat1 (blue) – weak signs of regeneration, Cat2 (green) – visible signs of development, and Cat3 (yellow) – complete and optimal regeneration. The y-axis represents the number of calli per quality category, and the x-axis the experimental conditions (control vs. TGF-β1) for each stage.

Figure 2b illustrates the qualitative scoring method used, with a series of images obtained during the regeneration process at three types of physiological tissue structures of the *Cervil* accession: cell clusters (day 30), buds (day 45) and isolated plants (day 60). The visual quality scores associated with each stage ranked from 0 (poor or no regeneration) to 3 (optimal regeneration). The + indicates whether the explant is supplemented with regulator TGF-β1. The highest scores (3) corresponded to well-developed plants with a morphology close to that expected for normal plants. Note that the majority of category 2 and 3 samples, whatever the stage, were obtained by supplementation with TGF-β1, unlike categories 0 and 1 which mainly came from the control without supplementation. These observations confirmed the positive effect of TGF-β1 on complete regeneration, promoting not only dedifferentiation but also the organization of tissues into differentiated cells and then the maturation of organs.

These observations are confirmed on Figure 2c with the results of the ratings averaged over three years of successive experiments. Supplementation with TGF-β1 significantly increased the frequency of category 3 calluses at all stages. At each stage, the frequency of category 0 (explant considered senescent) was reduced in favor of categories 2 and 3, while category 1 remained stable. Category 2 increased sharply at the beginning of the cell cluster stage while category 3 increased at the budding stage compared to the control without addition of TGF-β1. The counts per accession are presented in the Supplementary Table 6.

#### Effect of TGF-β1 on six accessions (years 2018 and 2019)

Figure 3 shows the regeneration frequencies of explants of six accessions (*Cervil, Criollo, Ferum, LA1420, Levovil* and *Stupike*) in response to TGF-β1 treatment, compared to a control condition without supplementation, at four stages of regeneration, over two experiments. The regeneration frequencies of different stages were improved by the addition of TGF-β1 for all accessions, but the improvement intensity depended on the genotype and stage. The counts per accession are presented in the Supplementary Table 7. The increase in cell clusters was statistically significant (p-value = 0.000284). However *Stupike* showed less improvement at the first stage (red arrow). Budding was also significantly improved by TGF-β1 (p-value = 2.17. 10^-08^). The budding frequency increased for the majority of accessions, although the magnitude of this increase varied. Accessions such as *Cervil, Criollo* and *Stupike* were particularly sensitive to the treatment.

**Figure 3:**
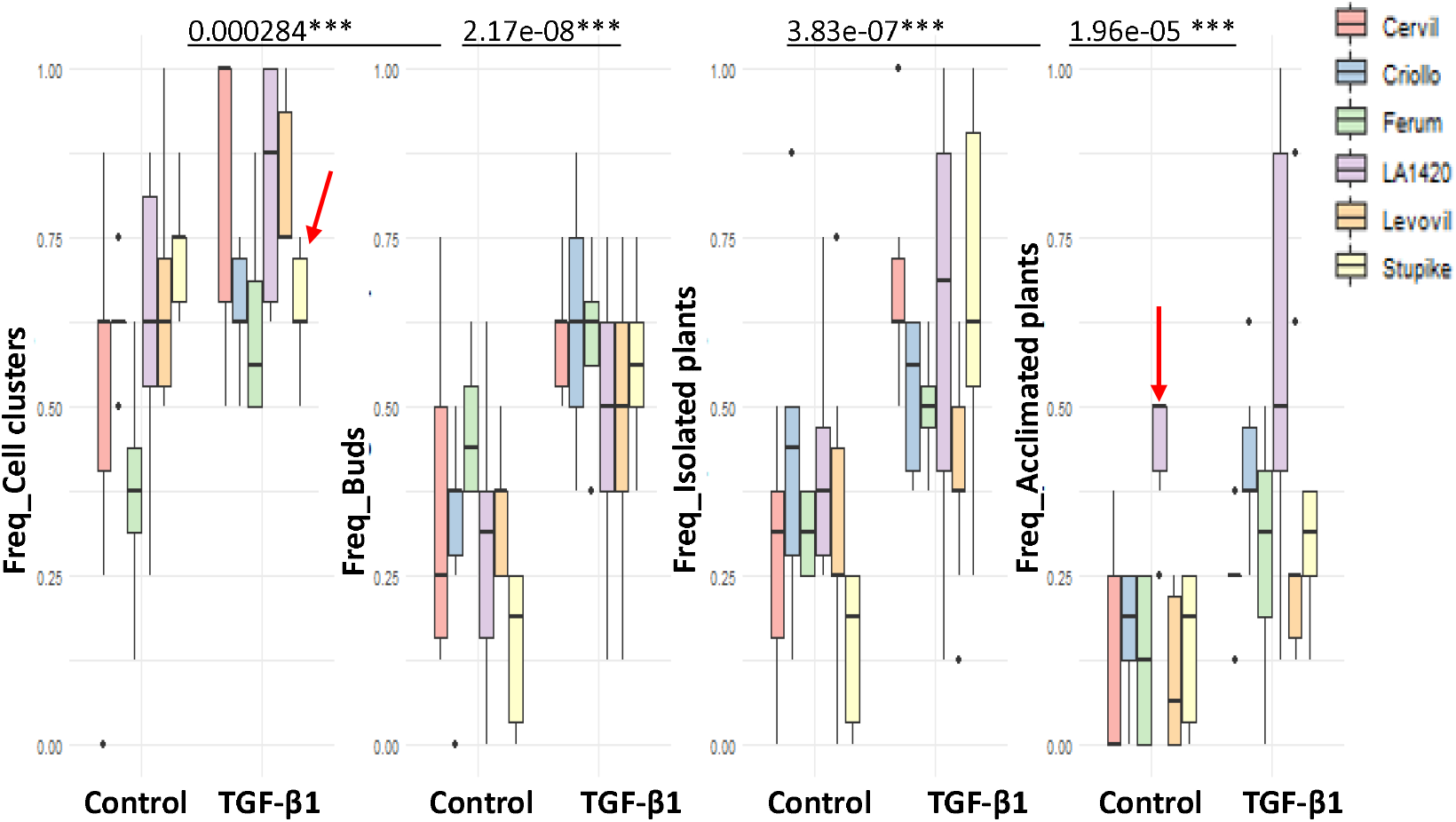
Comparison of regeneration frequencies for two consecutive years (2018 and 2019) on explants transformed for KO genes Mal30, FRK3, Lin5, Suc3 of six accessions (*Cervil, Criollo, Ferum, LA1420, Levovil* and *Stupike*) for two conditions, with TGF-β1 supplementation (2 mg/mL) compared to the control condition without supplementation, at four regeneration stages (cell clusters, buds, isolated seedlings and acclimated plants). Each colored bar represents a specific accession and the height of each bar indicates the regeneration frequency for that accession. The significance levels are 0.001 whatever the stage and condition.

The development of isolated plants followed the trend of the previous stages, with a significant increase in regeneration frequencies under the effect of TGF-β1 (p-value 3.83.10^-07^). However, the inter-genotype variability was more pronounced, with *Stupike* showing a better response than others at this stage.

At the stage of acclimatized plants, the trends were similar with an improvement under the effect of TGF-β1 (p-value = 1.96.10^-5^) although the impact seemed less pronounced at this stage compared to earlier stages, such as the less marked improvement of the non-sensitive accession *LA1420* (red arrow). The accession *Criollo*, very sensitive to acclimation, responded very well to TGF-β1.

We also observed a reduction in the physiological and behavioral imbalance of the accessions during the regeneration process across all genotypes under the influence of TGF-β1. Specifically, TGF-β1 supplementation enhanced the regeneration of transformed calli in the recalcitrant genotype *Cervil*, promoted regeneration in the low-regenerating genotype *Stupike*, and increased regeneration frequencies in transformable genotypes (*Ferum, Criollo, Levovil*, and *LA1420*). TGF-β1 promoted differentiation at the embryonic stage, except for *Stupike*, but improved it at later stages. The frequency of the three other stages—induction, development, and acclimation—was significantly higher compared to the control for all genotypes. Additionally, TGF-β1 supplementation improved greenhouse acclimation for the *Criollo* genotype.

Examining the results year by year, the effect of the growth factor TGF-β1 was significant at all stages for both 2018 and 2019 across all accessions (Supplementary Figure S5). In 2019, TGF-β1 had a significantly positive impact on all accessions except *LA1420* at every growth stage. For *Cervil*, TGF-β1 showed a consistently positive effect at all stages (Supplementary Figure S6,). Additionally, the impact of TGF-β1 on the quality of explants in culture was confirmed for all six accessions (Figure S7)

#### Visual effect of TGF-β1 on four accessions

Two series of pictures taken at 43 days (bud stage) and 55 days (isolated plant stage) after transformation illustrate the improvement in regeneration with TGF-β1 supplementation (Supplementary plate 1, 2). The effect of TGF-β1 promoted differentiation and cell multiplication of the accessions *Cervil, Stupike, Criollo* and *Levovil*. For *Cervil*, regeneration was weak or absent in the absence of TGF-β1 while with its addition, the response was linear and more pronounced in the later experiment, suggesting that the growth factor facilitated the initiation of budding and maintained its effect during later stages of development. In *Stupike*, regeneration was more pronounced with numerous and better-developed buds, indicating a progressive positive response to TGF-β1, which was reflected in the formation of numerous seedlings. In *Criollo*, the response to the growth regulator was visible from the bud stage, with significantly greater regeneration compared to the control. The *Levovil* accession exhibited lower regeneration both in the absence of TGF-β1 and compared to the *Criollo*. However, its response was notable at the seedling stage.

#### Gene expression of two transformed accessions obtained with the TGF-1β regulator

In order to confirm that the effect of the regulator TGF-β1 improved transformation and did not produce artifactual transgenic lines, we assessed gene expression on secondary mutants of *Cervil* and *Levovil* for three genes (Suc3: solyc03g083910, FRK3: solyc02g091490 and Lin5: solyc09g010080). The mutations caused on these three genes in the KO mutants of *Cervil* and *Levovil* systematically decreased the expression of these genes regardless of the accessions, confirming that transgenesis was functional (Supplementary Figures S8).

#### Comparison of the effect of three growth factor regulators, TGF-β1, EGF and IL-1β on plant regeneration

In 2022, we compared the regeneration frequencies at three stages with and without supplementation of two cytokines, the growth factors (TGF-β1 and EGF) and a pro-inflammatory paracrine factor (IL-1β) on *Cervil* accession. The cell cluster stage was significantly favored by the three factors although the effects observed at the bud and isolated plant stages were not significant. The pro-inflammatory regulator IL-1β had a slight advantage over the two others (p = 0.0001 vs p = 0.001) (Figure 4a). The results of the transgenesis experiment are presented in the supplemental data Table 8.

**Figure 4:**
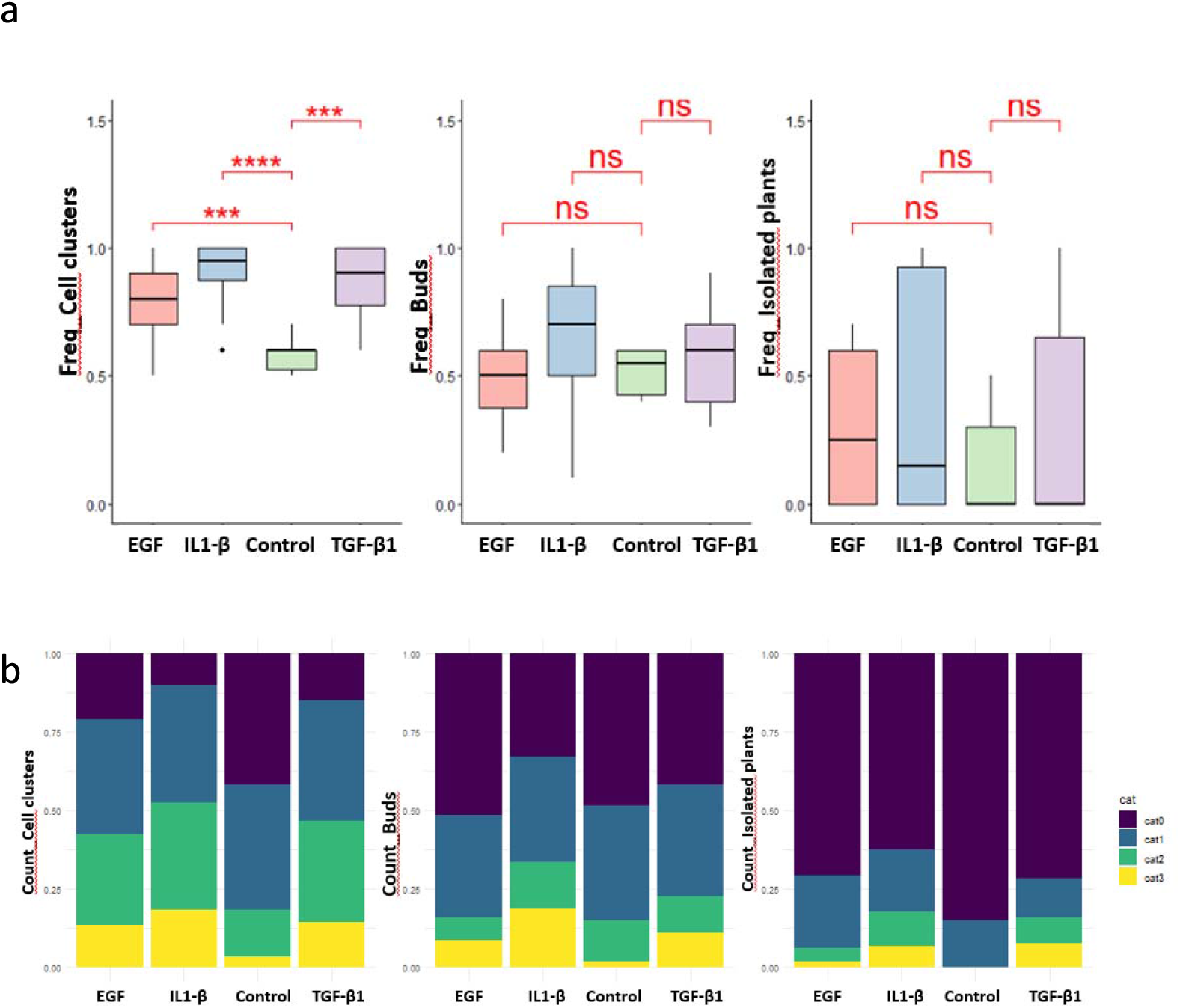
**a**. Comparison of the regeneration frequencies of explants transformed with the FRK3 - KI gene by the addition of three cytokine-type growth factors, TGF-β1 and EGF, and a pro-inflammatory factor of the interleukin 1β (IL) type, for 2 conditions with and without supplementation, for three stages (cell clusters, buds, isolated seedlings) for the year 2022 (12 values are analyzed per condition and 6 values for the control condition). The significance levels are ^***^ p= 0.001 and ^****^ p= 0.0001 and ‘NS’ for non-significant. **b**. Comparison of the number of transformed explants with and without supplementation of three growth factors (cytokine type: TGF-β1, EGF and paracrine pro-inflammatory type: interleukin 1β) on the *Cervil* genotype in 2022 for 3 stages (cell clusters, buds, isolated seedlings). The four quality scores are identical to those in Figure 2b. The Y axis indicates the number of calli for each regeneration quality category. The X axis indicates the experimental conditions for each developmental stage (cell clusters, buds, isolated plants).

The quality of the regeneration during the transformation process of *Cervil* was improved by the three growth regulators (Figure 4b). Categories 1, 2 and 3 were all increased at the cell cluster stage; the category 3 was more clearly increased at the budding stage with a slight advantage for IL-1β; the categories 2 and 3 increased at the isolated plant stage to the detriment of the categories 0 and 1 compared to the control. This improvement was illustrated in Supplementary Figure S9. The control condition (without supplementation of any regulator) showed inactive cotyledons and hypocotyls with a pale color of homogeneous texture, suggesting a vegetative arrest or an upcoming senescence. For TGF-β1, we observed clear signs of more pronounced regeneration with the appearance of green structures and the emergence of denser areas and potential buds indicating a positive response of the regulator in both tissues; for the EGF regulator, the explants also showed better regeneration, with dark green areas characteristic of early development, as for the IL-1β response, where we observed a more limited cell proliferation but typical of a future cell cluster.

## Discussion

Numerous studies have emphasized the variation in regeneration capacity among tomato accessions during the transgenesis process, spanning from the transformation phase to the production of mutant plants (Fuentes et al. 2008). These variations lead to distinct physiological responses at each stage of transgenesis, exposing the unique vulnerabilities of each accession. Growth bottlenecks associated with transgenesis stress can result from several factors, including the unbalanced activation of the innate immune response induced by the introduced gene, the transformation process itself, and the disruption of phytohormone levels linked to plant development and the production of reactive oxygen species (ROS) (Aurora and Olson 2014; Heyman et al. 2018; Abnave and Ghigo 2019).

### Role of plant kinase receptors

The balance between the complex mechanisms of repression and activation of specific plant kinases within the receptor tyrosine kinase (RTK) family has been extensively studied (Belda-Palazón et al. 2020). In tomato, a connection has been established between the cytoplasmic kinase receptor TRK1 and several key processes, including chitin-induced fungal resistance, reactive oxygen species (ROS) accumulation, and the expression of genes involved in immune responses (Jaiswal et al. 2022). Receptor tyrosine kinases, related to the regulators EGF and TGF-β1, share similarities with the animal immune system and have been well characterized in Arabidopsis thaliana (Shiu and Bleecker 2001; Jose et al. 2020).

Plants use immune receptors homologous to those of mammals to detect infections and activate their defenses. Among them, LRR receptors, notably RLK and RLP, play a key role in innate immunity by perceiving various danger signals. The more abundant LRR-RLKs regulate essential processes such as development and stress response. Their study, particularly in tomato and Arabidopsis, sheds light on their network organization and opens up perspectives for better understanding plant resistance to pathogens (Zhou and Zhang 2020) and the defense mechanisms involved during the transgenesis process.

Homology analysis revealed some similarities between mammalian and tomato protein sequences with scores indicating a high probability of non-coincidental similarity. Despite the presence of some gaps in the alignments that could be explained by the distance between the organisms, these matches may correspond to common functional and evolutionary implications. Moreover, the BLAST results between mammalian and tomato receptors show a significant conservation of functional domains, notably kinase domains, despite their divergent evolution. Tomato interacting protein 1 (solyc12g098990, Figure S2d) is homologous to a serine threonine protein kinase that is involved in the signaling cascade of the immune response linked to the murine EGF receptor is published by (Zhou et al. 1995).

### Ligands as regulators of regeneration

In mammals, ligands such as TGF-β1 and EGF are long-lived, highly diluted growth factors that interact systemically in the general circulation and exert their effects by binding to their receptors, which themselves have highly diverse extracellular domains, thereby activating networked signaling cascades. In contrast, the proinflammatory cytokine IL-1β acts in a juxtacrine manner, influencing the nearby environment and providing contact immune response properties during transformation. It signals via two receptors, IL-1R1 and IL-1R2, triggering the production of other proinflammatory cytokines in target cells and initiating acute-phase responses to infection during injuries to explant cuts (Combarnous 2013).

Plants use immune receptors similar to those of mammals to detect infections and coordinate their defense responses, as explored by Zhou and Zhang (2020).

LRR receptors, key to innate immunity in mammals, share strong homology with plant RLKs and RLPs, which provide early defense by detecting various pathogens, including lipopolysaccharides (LPS) from Gram-negative bacteria during transgenesis (Clos 2012). Among them, histidine receptor kinases (LRR-RLKs), related to interleukin regulators, represent the largest group of cell surface receptors in plants, including somatic SERK kinases, which play key roles in human embryonic development and plant immunity (Walker 1994; Chevalier and Walker 2005; Hosseini et al. 2020). This receptor family perceives various biotic and abiotic stimuli, influencing key biological processes, particularly described in plants such as tomato and Arabidopsis (Wei et al. 2015; Cheng et al. 2021; Soltabayeva et al. 2022).

Our data revealed that supplying human and murine growth factors enhanced both the quality of regeneration and the transformation rate of explants from various tomato accessions, each transformed with different CRISPR-Cas9 constructs. Each ligand positively impacted the regeneration of embryonic cells.

Following our results, it could be interesting to synthesize plant ligands that precisely adapt to plant receptors to reactivate signaling and communication pathways disrupted by DNA breaks (for example during CRISPR transgenesis) and complex transgenesis processes, sources growth blockages in transformed plants. Phytocytokines appear capable of influencing the expression of genes involved in kinases and immune responses, thus playing a role in the process of cellular regeneration and reprogramming. Current research is focusing on the use of peptides capable of activating WIND1, a key regulator of injury-induced cellular reprogramming, by binding to receptors such as kinase 1 (Yang et al. 2024). Thus, the use of growth regulators such as “phytocytokines and pro-inflammatory molecules” to improve regeneration rates in plant transgenesis is a useful tool. It is time to consider synthesizing ligands from these superfamilies to boost regeneration in recalcitrant accessions. It could be relevant to use a pro-inflammatory factor, such as an interleukin, to facilitate dedifferentiation during co-culture by promoting a balanced immune response. This could be combined with a long-acting growth factor like TGF-β1 or EGF to stimulate cell differentiation and division.

### Cell environment

Today, a functional link between the variability of gene expression and the process of cellular differentiation has been demonstrated. The increase in gene variability is accompanied by an acceleration of cell differentiation (Guillemin et al. 2019). The more the sources of gene variability increase, at the time of coculture, the greater the chances of still undifferentiated cells to form clusters and produce structured and homogeneous tissues.

Variability in gene expression is therefore an essential component of the differentiation mechanism. Therefore, the current issue to improve the quality and quantity of transformation would be to improve our understanding of the functional interdependence between gene expression variability and the differentiation process at early stages of the transformation process.

## Conclusion

Genetic transformation represents a delicate balance between the aggressiveness of Agrobacterium and the initiation of embryogenesis, following a series of mechanisms involving MAP kinases. Recent research has shown that enhancing the plant transformation process depends on a deeper understanding of embryo development during cellular repair, which is closely linked to the immune response during infection.

The introduction of growth regulators from human and murine systems has emerged as a promising strategy to overcome the regeneration challenges faced by recalcitrant accessions. These results open up interesting possibilities for optimizing genetic transformation protocols, particularly for accessions presenting regeneration difficulties. This strategy could also be extended to other plant species with similar regeneration profiles, thereby contributing to the advancement of plant biotechnology.

## Supplementary materials

**Supplementary Figure S1:**

Protein BLAST results using the human receptor sequence P36897 TGFR1_HUMAN as a query against *Solanum lycopersicum*-specific protein sequence databases

**Supplementary Figure S2:**

Protein BLAST results using the murine receptor sequence Q01279 EGFR_MURIN as a query against *Solanum lycopersicum*-specific protein sequence databases

**Supplementary Figure S3:**

Protein BLAST results using the murine receptor sequence P13504 IL1RI_MURIN as a query against *Solanum lycopersicum*-specific protein sequence databases

**Supplementary Figure S4:** Potein BLAST using the murine receptor sequence P27931 IL1RII_MURIN as query against *Solanum lycopersicum*-specific protein sequence databases

**Supplementary Figure S5:** Comparison of the relative frequencies of the different stages of regeneration (cell clusters, buds, isolated seedlings and plants acclimated in a greenhouse) over two consecutive years (2018 and 2019) of transformed explants of 6 accessions (*Cervil*, Levovil, Ferum, Stupike, LA1420 and Criollo) for 2 conditions, with supplementation of TGF-β1 at a concentration of 2 mg/mL (TGF-β1) compared to the control condition without supplementation.

**Supplementary Figure S6:** Comparison of the regeneration frequency in 2019 on transformed explants for 2 conditions, with TGF-β1 supplementation compared to the control condition without supplementation at 4 stages

**Supplementary Figure S7:** Comparison of the quality of explants transformed into regeneration in 2019 for two conditions, with supplementation of TGF-β1 compared to the control condition without supplementation, for 3 stages (cell cluster, buds, isolated plantlets).

**Supplementary Figure S8:** Comparison of the effect of TGF-β1 vs. control for the *Cervil* and *Stupike* accessions.

**Supplementary Figure S9:** Comparison of the effect of TGF-β1 vs. control for the *Criollo* and *Levovil* accessions.

**Supplementary Figure S10:** Gene expression in edited and control plants

**Supplementary Figure S11:** Illustration of the effect of three growth regulators TGF-β1, EGF and IL-1β on the quality of transformed explants of hypocotyls and cotyledons regenerating at the cell cluster stage of the *Cervil* accession.

**Supplementary Table 1 :** details of constructs

**Supplementary Table 2 :** Map of plasmids used for construct cloning

**Supplementary Table 3 :** Protein sequences of the regulators

**Supplementary Table 4 :** Composition of media used for the culture of agrobacteria and the in vitro culture of calluses and transgenic plants

**Supplementary Table 5**: Genome editing results of secondary transformants of *Cervil* and *Levovil* accessions obtained with the addition of TGF during transgenesis for the target genes Suc3, FRK3 and Lin5

**Supplementary Table 6 :** Transgenesis experiment 2018 results (number of explants, cell clusters and plants at each step)

**Supplementary Table 7 :** Transgenesis experiment 2019 (number of explants, cell clusters and plants at each step)

**Supplementary Table 8 :** Transgenesis experiment 2022 (number of explants, cell clusters and plants at each step)

## Acknowledgement

We are thankful to the greenhouse staff of UE A2M “Arboriculture et Maraichage Méditerranéens” for taking care of the plant infrastructures and to CRBLeg for the conservation and access to genetic resources.

## Author contribution

CG: experiment conceptualization; AG, JG, EP : data acquisition; JB : statistical analysis; MC and CG: writing the manuscript

## Conflict of Interest

The authors declare that they have no conflict of interest.

## Notes

### Competing Interest Statement

The authors have declared no competing interest.

### Funding

- ANR-24-PESV-0001

## References

Abnave P, Ghigo E (2019) Role of the immune system in regeneration and its dynamic interplay with adult stem cells. Seminars in Cell & Developmental Biology 87:160–168. 10.1016/j.semcdb.2018.04.002

Albert E, Duboscq R, Latreille M, et al (2018) Allele-specific expression and genetic determinants of transcriptomic variations in response to mild water deficit in tomato. The Plant Journal 96:635–650. 10.1111/tpj.14057

Aurora AB, Olson EN (2014) Immune Modulation of Stem Cells and Regeneration. Cell Stem Cell 15:14–25. 10.1016/j.stem.2014.06.009

Banno H, Ikeda Y, Niu Q-W, Chua N-H (2001) Overexpression of Arabidopsis ESR1 Induces Initiation of Shoot Regeneration. Plant Cell 13:2609–2618. 10.1105/tpc.010234

Bashyal S, Gautam CK, Müller LM (2024) CLAVATA signaling in plant-environment interactions. Plant Physiol 194:1336–1357. 10.1093/plphys/kiad591

Bauchet G, Grenier S, Samson N, et al (2017) Identification of major loci and genomic regions controlling acid and volatile content in tomato fruit: implications for flavor improvement. New Phytol 215:624–641. 10.1111/nph.14615

Belda-Palazón B, Adamo M, Valerio C, et al (2020) A dual function of SnRK2 kinases in the regulation of SnRK1 and plant growth. Nat Plants 6:1345–1353. 10.1038/s41477-020-00778-w

Bergey DR, Howe GA, Ryan CA (1996) Polypeptide signaling for plant defensive genes exhibits analogies to defense signaling in animals. Proceedings of the National Academy of Sciences 93:12053–12058. 10.1073/pnas.93.22.12053

Brooks C, Nekrasov V, Lippman ZB, Van Eck J (2014) Efficient Gene Editing in Tomato in the First Generation Using the Clustered Regularly Interspaced Short Palindromic Repeats/CRISPR-Associated9 System. Plant Physiology 166:1292–1297. 10.1104/pp.114.247577

Bull T, Debernardi J, Reeves M, et al (2023) GRF–GIF chimeric proteins enhance in vitro regeneration and Agrobacterium-mediated transformation efficiencies of lettuce (Lactuca spp.). Plant Cell Rep 42:629–643. 10.1007/s00299-023-02980-4

Carpenter G, Liao H-J (2009) Trafficking of receptor tyrosine kinases to the nucleus. Exp Cell Res 315:1556–1566. 10.1016/j.yexcr.2008.09.027

Causse M, Bénéjam J, Bineau E, et al (2022) Genetic control of tomato fruit quality: from QTL mapping to Genome Wide Association studies and breeding. Comptes Rendus Biologies 345:3–13. 10.5802/crbiol.99

Cheng W, Wang Z, Xu F, et al (2021) Genome-Wide Identification of LRR-RLK Family in Saccharum and Expression Analysis in Response to Biotic and Abiotic Stress. Curr Issues Mol Biol 43:1632–1651. 10.3390/cimb43030116

Chetelat RT, Deverna JW, Bennett AB (1995) Introgression into tomato (Lycopersicon esculentum) of the L. chmielewskii sucrose accumulator gene (sucr) controlling fruit sugar composition. Theor Appl Genet 91:327–333. 10.1007/BF00220895

Chevalier D, Walker JC (2005) Functional genomics of protein kinases in plants. Brief Funct Genomic Proteomic 3:362–371. 10.1093/bfgp/3.4.362

Clos J (2012) L’immunité chez les animaux et les végétaux. Lavoisier

Combarnous Y (2013) Communications et signalisations cellulaires, 4ème. Lavoisier, Paris

Debernardi JM, Tricoli DM, Ercoli MF, et al (2020) A chimera including a GROWTH-REGULATING FACTOR (GRF) and its cofactor GRF-INTERACTING FACTOR (GIF) increases transgenic plant regeneration efficiency. 2020.08.23.263905

Dhugga KS (2022) Gene Editing to Accelerate Crop Breeding. Frontiers in Plant Science 13:

Doudna JA, Charpentier E (2014) The new frontier of genome engineering with CRISPR-Cas9. Science 346:1258096. 10.1126/science.1258096

Doudna JA, Jinek M, Chylinski K, Charpentier E (2018) Methods and compositions for RNA-directed target DNA modification and for RNA-directed modulation of transcription. 32

Fan L, Li R, Pan J, et al (2015) Endocytosis and its regulation in plants. Trends Plant Sci 20:388–397. 10.1016/j.tplants.2015.03.014

Fuentes AD, Ramos PL, Sánchez Y, et al (2008) A transformation procedure for recalcitrant tomato by addressing transgenic plant-recovery limiting factors. Biotechnology Journal 3:1088–1093. 10.1002/biot.200700187

Gao L, Gonda I, Sun H, et al (2019) The tomato pan-genome uncovers new genes and a rare allele regulating fruit flavor. Nat Genet 51:1044–1051. 10.1038/s41588-019-0410-2

Garchery C, Gest N, Do PT, et al (2013) A diminution in ascorbate oxidase activity affects carbon allocation and improves yield in tomato under water deficit. Plant, Cell & Environment 36:159–175. 10.1111/j.1365-3040.2012.02564.x

Gerszberg A, Hnatuszko-Konka K, Kowalczyk T, Kononowicz AK (2015) Tomato (Solanum lycopersicum L.) in the service of biotechnology. Plant Cell Tiss Organ Cult 120:881–902. 10.1007/s11240-014-0664-4

Gostimskaya I (2022) CRISPR–Cas9: A History of Its Discovery and Ethical Considerations of Its Use in Genome Editing. Biochemistry (Mosc) 87:777–788. 10.1134/S0006297922080090

Guillemin A, Duchesne R, Crauste F, et al (2019) Drugs modulating stochastic gene expression affect the erythroid differentiation process. PLOS ONE 14:e0225166. 10.1371/journal.pone.0225166

Gust AA, Pruitt R, Nürnberger T (2017) Sensing Danger: Key to Activating Plant Immunity. Trends in Plant Science 22:779–791. 10.1016/j.tplants.2017.07.005

Hamza S, Chupeau Y (1993) Re-evaluation of Conditions for Plant Regeneration and Agrobacterium-Mediated Transformation from Tomato (Lycopersicon esculentum). Journal of Experimental Botany 44:1837–1845. 10.1093/jxb/44.12.1837

Henri⍰Desaint A, re⍰Héreil JB-M, Yol e C, et al Integration of QTL and transcriptome approaches for the identification of genes involved in tomato response to nitrogen deficiency

Héreil A, Guillaume M, Duboscq R, et al Characterisation of a major QTL for sodium accumulation in tomato grown in high salinity. Plant, Cell & Environment n/a: 10.1111/pce.15082

Heyman J, Canher B, Bisht A, et al (2018) Emerging role of the plant ERF transcription factors in coordinating wound defense responses and repair. Journal of Cell Science 131:jcs208215. 10.1242/jcs.208215

Hosseini S, Schmidt EDL, Bakker FT (2020) Leucine-rich repeat receptor-like kinase II phylogenetics reveals five main clades throughout the plant kingdom. Plant J 103:547–560. 10.1111/tpj.14749

Ichihashi Y, Hakoyama T, Iwase A, et al (2020) Common Mechanisms of Developmental Reprogramming in Plants—Lessons From Regeneration, Symbiosis, and Parasitism. Front Plant Sci 11:1084. 10.3389/fpls.2020.01084

Ikeuchi M, Favero DS, Sakamoto Y, et al (2019) Molecular Mechanisms of Plant Regeneration. Annu Rev Plant Biol 70:377–406. 10.1146/annurev-arplant-050718-100434

Ikeuchi M, Ogawa Y, Iwase A, Sugimoto K (2016) Plant regeneration: cellular origins and molecular mechanisms. Development 143:1442–1451. 10.1242/dev.134668

Imai R, Hamada H, Liu Y, et al (2020) In planta particle bombardment (iPB): A new method for plant transformation and genome editing. Plant Biotechnol (Tokyo) 37:171–176. 10.5511/plantbiotechnology.20.0206a

Iwase A, Mita K, Nonaka S, et al (2015) WIND1-based acquisition of regeneration competency in Arabidopsis and rapeseed. J Plant Res 128:389–397. 10.1007/s10265-015-0714-y

Jaiswal N, Liao C-J, Mengesha B, et al (2022) Regulation of plant immunity and growth by tomato receptor-like cytoplasmic kinase TRK1. New Phytologist 233:458–478. 10.1111/nph.17801

Jose J, Ghantasala S, Roy Choudhury S (2020) Arabidopsis Transmembrane Receptor-Like Kinases (RLKs): A Bridge between Extracellular Signal and Intracellular Regulatory Machinery. Int J Mol Sci 21:4000. 10.3390/ijms21114000

Kareem A, Radhakrishnan D, Wang X, et al (2016) Protocol: a method to study the direct reprogramming of lateral root primordia to fertile shoots. Plant Methods 12:27. 10.1186/s13007-016-0127-5

Kausch AP, Nelson-Vasilchik K, Hague J, et al (2019) Edit at will: Genotype independent plant transformation in the era of advanced genomics and genome editing. Plant Sci 281:186–205. 10.1016/j.plantsci.2019.01.006

Kong J, Martin-Ortigosa S, Finer J, et al (2020) Overexpression of the Transcription Factor GROWTH-REGULATING FACTOR5 Improves Transformation of Dicot and Monocot Species. Front Plant Sci 11:572319. 10.3389/fpls.2020.572319

Long Y, Yang Y, Pan G, Shen Y (2022) New Insights Into Tissue Culture Plant-Regeneration Mechanisms. Frontiers in Plant Science 13:

Lor VS, Starker CG, Voytas DF, et al (2014) Targeted Mutagenesis of the Tomato PROCERA Gene Using Transcription Activator-Like Effector Nucleases. Plant Physiology 166:1288–1291. 10.1104/pp.114.247593

Lu Y, Meng Y, Zeng J, et al (2020) Coordination between GROWTH-REGULATING FACTOR1 and GRF-INTERACTING FACTOR1 plays a key role in regulating leaf growth in rice. BMC Plant Biology 20:200. 10.1186/s12870-020-02417-0

Maren NA, Duan H, Da K, et al (2022) Genotype-independent plant transformation. Horticulture Research 9:uhac047. 10.1093/hr/uhac047

Méndez-Hernández HA, Ledezma-Rodríguez M, Avilez-Montalvo RN, et al (2019) Signaling Overview of Plant Somatic Embryogenesis. Frontiers in Plant Science 10:

Minoia S, Petrozza A, D’Onofrio O, et al (2010) A new mutant genetic resource for tomato crop improvement by TILLING technology. BMC Res Notes 3:69. 10.1186/1756-0500-3-69

Montoya T, Nomura T, Farrar K, et al (2002) Cloning the tomato curl3 gene highlights the putative dual role of the leucine-rich repeat receptor kinase tBRI1/SR160 in plant steroid hormone and peptide hormone signaling. Plant Cell 14:3163–3176. 10.1105/tpc.006379

Nie S, Huang S, Wang S, et al (2019) Enhanced brassinosteroid signaling intensity via SlBRI1 overexpression negatively regulates drought resistance in a manner opposite of that via exogenous BR application in tomato. Plant Physiol Biochem 138:36–47. 10.1016/j.plaphy.2019.02.014

Pascual L, Xu J, Biais B, et al (2013) Deciphering genetic diversity and inheritance of tomato fruit weight and composition through a systems biology approach. J Exp Bot 64:5737–5752. 10.1093/jxb/ert349

Qiu D, Diretto G, Tavarza R, Giuliano G (2007) Improved protocol for Agrobacterium mediated transformation of tomato and production of transgenic plants containing carotenoid biosynthetic gene CsZCD. Scientia Horticulturae 112:172–175. 10.1016/j.scienta.2006.12.015

Rasheed A, Gill RA, Hassan MU, et al (2021) A Critical Review: Recent Advancements in the Use of CRISPR/Cas9 Technology to Enhance Crops and Alleviate Global Food Crises. Curr Issues Mol Biol 43:1950–1976. 10.3390/cimb43030135

Rehli M (2002) Of mice and men: species variations of Toll-like receptor expression. Trends in Immunology 23:375–378. 10.1016/S1471-4906(02)02259-7

Rothan C, Diouf I, Causse M (2019) Trait discovery and editing in tomato. The Plant Journal 97:73–90. 10.1111/tpj.14152

Ryan CA (2000) The systemin signaling pathway: differential activation of plant defensive genes. Biochimica et Biophysica Acta (BBA) - Protein Structure and Molecular Enzymology 1477:112–121. 10.1016/S0167-4838(99)00269-1

Ryan CA, Pearce G (1998) SYSTEMIN: A Polypeptide Signal for Plant Defensive Genes. Annual Review of Cell and Developmental Biology 14:1–17. 10.1146/annurev.cellbio.14.1.1

Sauvage C, Segura V, Bauchet G, et al (2014) Genome-Wide Association in Tomato Reveals 44 Candidate Loci for Fruit Metabolic Traits. Plant Physiology 165:1120. 10.1104/pp.114.241521

Schouten HJ, Tikunov Y, Verkerke W, et al (2019) Breeding Has Increased the Diversity of Cultivated Tomato in The Netherlands. Front Plant Sci 10:1606. 10.3389/fpls.2019.01606

Shiu S-H, Bleecker AB (2001) Receptor-like kinases from Arabidopsis form a monophyletic gene family related to animal receptor kinases. Proceedings of the National Academy of Sciences 98:10763–10768. 10.1073/pnas.181141598

Soltabayeva A, Dauletova N, Serik S, et al (2022) Receptor-like Kinases (LRR-RLKs) in Response of Plants to Biotic and Abiotic Stresses. Plants 11:2660. 10.3390/plants11192660

Sree Vidya CS, Manoharan M, Ranjit Kumar CT, et al (2000) Agrobacterium-mediated Transformation of Tomato (Lycopersicon esculentum var. Pusa Ruby) with Coat-protein Gene of Physalis Mottle Tymovirus. Journal of Plant Physiology 156:106–110. 10.1016/S0176-1617(00)80279-5

Stevens R, Buret M, Duffé P, et al (2007) Candidate Genes and Quantitative Trait Loci Affecting Fruit Ascorbic Acid Content in Three Tomato Populations. Plant physiology 143:1943–53. 10.1104/pp.106.091413

Sugimoto K, Temman H, Kadokura S, Matsunaga S (2019) To regenerate or not to regenerate: factors that drive plant regeneration. Current Opinion in Plant Biology 47:138–150. 10.1016/j.pbi.2018.12.002

Sun H-J, Uchii S, Watanabe S, Ezura H (2006) A Highly Efficient Transformation Protocol for Micro-Tom, a Model Cultivar for Tomato Functional Genomics. Plant and Cell Physiology 47:426–431. 10.1093/pcp/pci251

Tanksley SD (2004) The genetic, developmental, and molecular bases of fruit size and shape variation in tomato. Plant Cell 16 Suppl:S181–189. 10.1105/tpc.018119

The Tomato Genome Consortium (2012) The tomato genome sequence provides insights into fleshy fruit evolution. Nature 485:635–641. 10.1038/nature11119

Tiwari JK, Singh AK, Behera TK (2023) CRISPR/Cas genome editing in tomato improvement: Advances and applications. Front Plant Sci 14:1121209. 10.3389/fpls.2023.1121209

Truffault V, Riqueau G, Garchery C, et al (2018) Is monodehydroascorbate reductase activity in leaf tissue critical for the maintenance of yield in tomato? Journal of Plant Physiology 222:1–8. 10.1016/j.jplph.2017.12.012

Vats S, Kumar V, Mandlik R, et al (2023a) Reference Guided De Novo Genome Assembly of Transformation Pliable Solanum lycopersicum cv. Pusa Ruby. Genes 14:570. 10.3390/genes14030570

Vats S, Shivaraj SM, Sonah H, et al (2023b) Efficient Regeneration and Agrobacterium-Mediated Transformation Method For Cultivated and Wild Tomato. Plant Molecular Biology Reporter. 10.1007/s11105-023-01374-w

Walker JC (1994) Structure and function of the receptor-like protein kinases of higher plants. Plant Mol Biol 26:1599–1609. 10.1007/BF00016492

Waltz E (2021) GABA-enriched tomato is first CRISPR-edited food to enter market. Nature Biotechnology 40:9–11. 10.1038/d41587-021-00026-2

Wang S, Liu J, Zhao T, et al (2019) Modification of Threonine-1050 of SlBRI1 regulates BR Signalling and increases fruit yield of tomato. BMC Plant Biology 19:256. 10.1186/s12870-019-1869-9

Wang S, Lv S, Zhao T, et al (2021) Modification of Threonine-825 of SlBRI1 Enlarges Cell Size to Enhance Fruit Yield by Regulating the Cooperation of BR-GA Signaling in Tomato. Int J Mol Sci 22:7673. 10.3390/ijms22147673

Wang P, Si H, Li C, Xu Z, Guo H, Jin S, Cheng H. Plant genetic transformation: achievements, current status and future prospects. Plant Biotechnol J. 2025 Jun;23(6):2034–2058. doi: 10.1111/pbi.70028.

Wei Z, Wang J, Yang S, Song Y (2015) Identification and expression analysis of the LRR-RLK gene family in tomato (Solanum lycopersicum) Heinz 1706. Genome 58:121–134. 10.1139/gen-2015-0035

Yang W, Zhai H, Wu F, et al (2024) Peptide REF1 is a local wound signal promoting plant regeneration. Cell 187:3024-3038.e14. 10.1016/j.cell.2024.04.040

Ye J, Wang X, Hu T, et al (2017) An InDel in the Promoter of Al-ACTIVATED MALATE TRANSPORTER9 Selected during Tomato Domestication Determines Fruit Malate Contents and Aluminum Tolerance. The Plant Cell 29:2249–2268. 10.1105/tpc.17.00211

Zanor MI, Osorio S, Nunes-Nesi A, et al (2009) RNA interference of LIN5 in tomato confirms its role in controlling Brix content, uncovers the influence of sugars on the levels of fruit hormones, and demonstrates the importance of sucrose cleavage for normal fruit development and fertility. Plant Physiol 150:1204–1218. 10.1104/pp.109.136598

Zhou J, Loh YT, Bressan RA, Martin GB (1995) The tomato gene Pti1 encodes a serine/threonine kinase that is phosphorylated by Pto and is involved in the hypersensitive response. Cell 83:925–935. 10.1016/0092-8674(95)90208-2

Zhou J-M, Zhang Y (2020) Plant Immunity: Danger Perception and Signaling. Cell 181:978–989. 10.1016/j.cell.2020.04.028

