## supplementary figures for "Mammalian growth-regulating factors enhance regeneration of recalcitrant transgenic tomato accessions"

### Slide 1
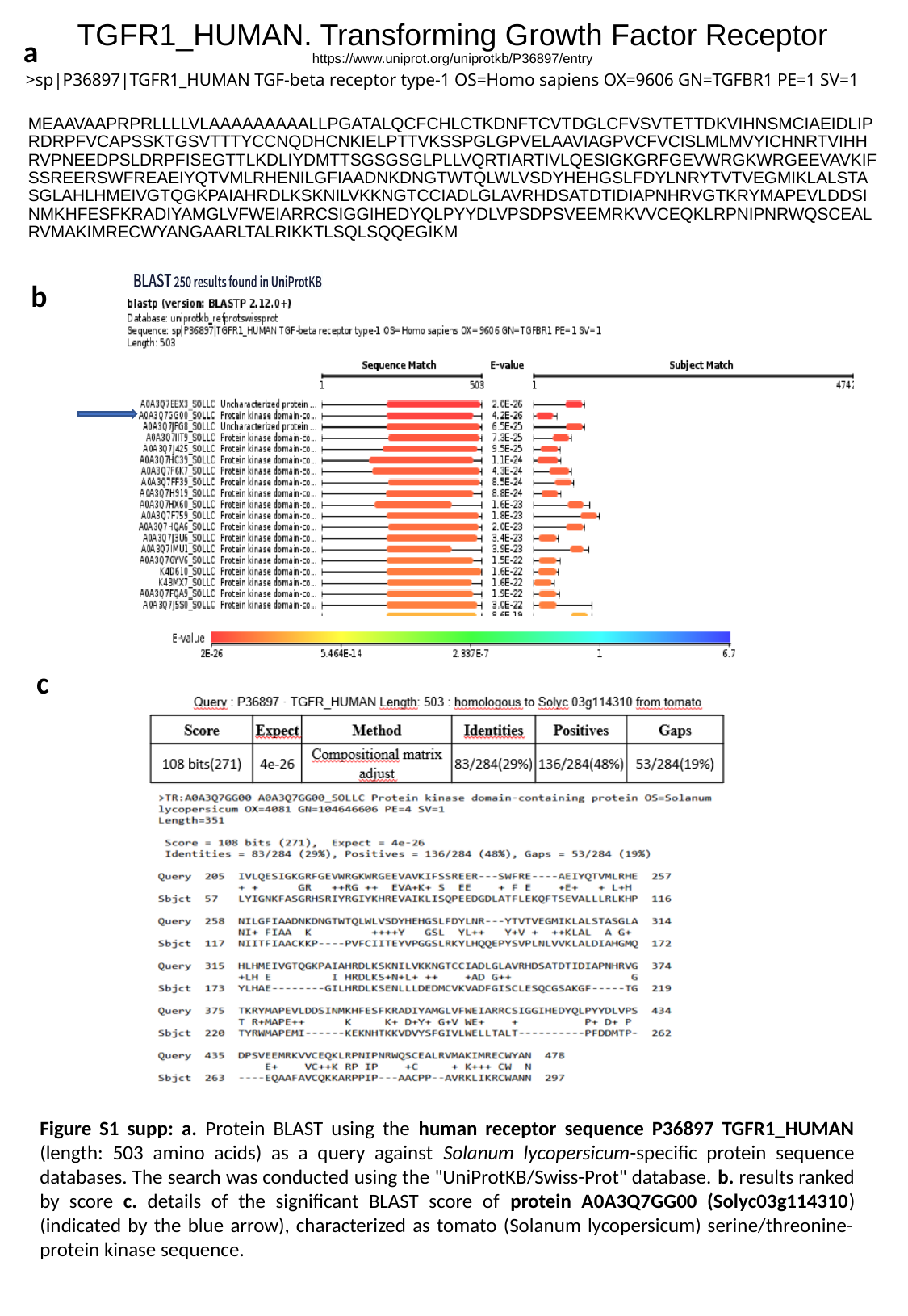

a
>sp|P36897|TGFR1_HUMAN TGF-beta receptor type-1 OS=Homo sapiens OX=9606 GN=TGFBR1 PE=1 SV=1
TGFR1_HUMAN. Transforming Growth Factor Receptor
https://www.uniprot.org/uniprotkb/P36897/entry
MEAAVAAPRPRLLLLVLAAAAAAAAALLPGATALQCFCHLCTKDNFTCVTDGLCFVSVTETTDKVIHNSMCIAEIDLIPRDRPFVCAPSSKTGSVTTTYCCNQDHCNKIELPTTVKSSPGLGPVELAAVIAGPVCFVCISLMLMVYICHNRTVIHHRVPNEEDPSLDRPFISEGTTLKDLIYDMTTSGSGSGLPLLVQRTIARTIVLQESIGKGRFGEVWRGKWRGEEVAVKIFSSREERSWFREAEIYQTVMLRHENILGFIAADNKDNGTWTQLWLVSDYHEHGSLFDYLNRYTVTVEGMIKLALSTASGLAHLHMEIVGTQGKPAIAHRDLKSKNILVKKNGTCCIADLGLAVRHDSATDTIDIAPNHRVGTKRYMAPEVLDDSINMKHFESFKRADIYAMGLVFWEIARRCSIGGIHEDYQLPYYDLVPSDPSVEEMRKVVCEQKLRPNIPNRWQSCEALRVMAKIMRECWYANGAARLTALRIKKTLSQLSQQEGIKM
b
c
Figure S1 supp: a. Protein BLAST using the human receptor sequence P36897 TGFR1_HUMAN (length: 503 amino acids) as a query against Solanum lycopersicum-specific protein sequence databases. The search was conducted using the "UniProtKB/Swiss-Prot" database. b. results ranked by score c. details of the significant BLAST score of protein A0A3Q7GG00 (Solyc03g114310) (indicated by the blue arrow), characterized as tomato (Solanum lycopersicum) serine/threonine-protein kinase sequence.

### Slide 2
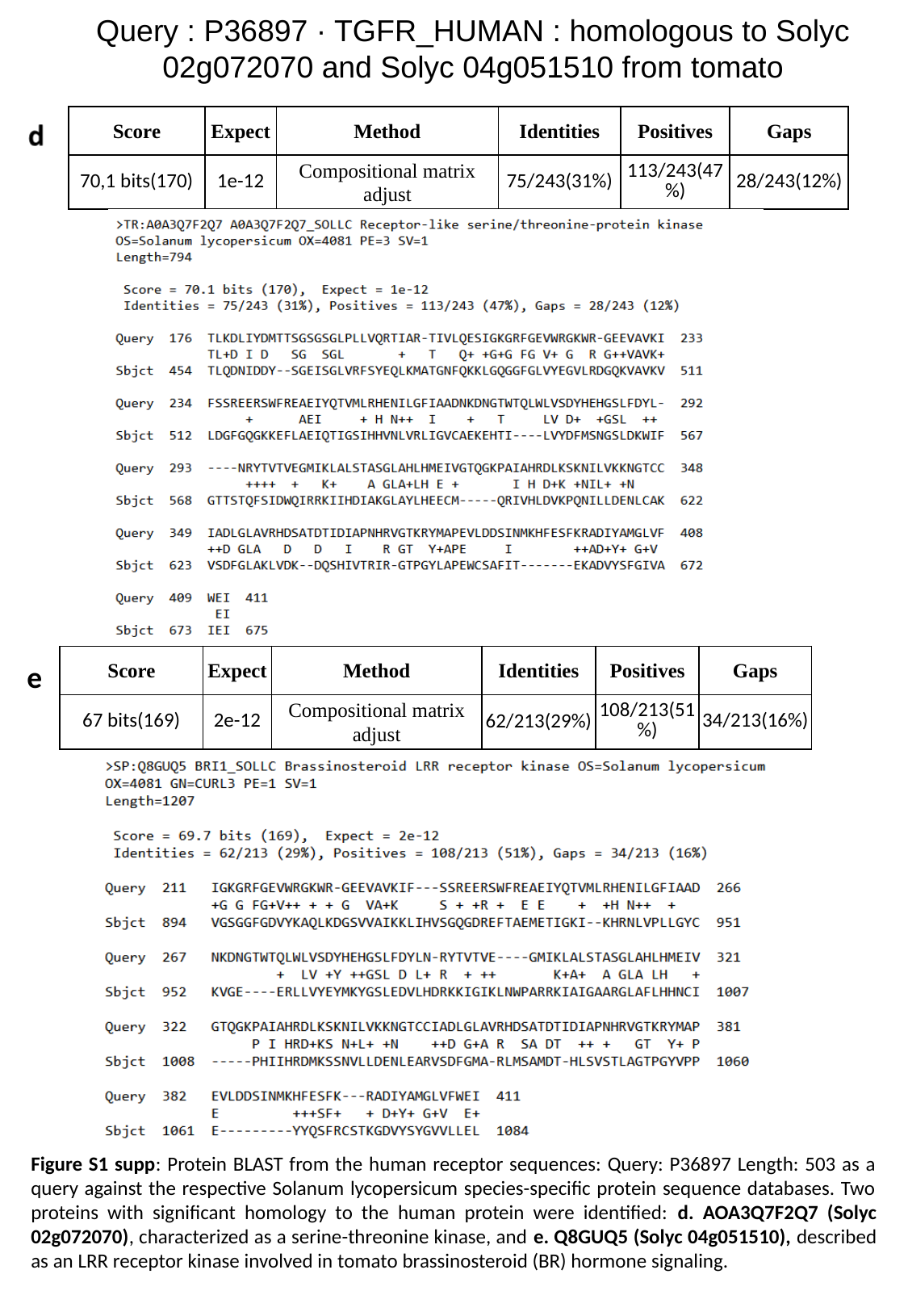

Query : P36897 · TGFR_HUMAN : homologous to Solyc 02g072070 and Solyc 04g051510 from tomato
| | | | | | |
| --- | --- | --- | --- | --- | --- |
| Score | Expect | Method | Identities | Positives | Gaps |
| 70,1 bits(170) | 1e-12 | Compositional matrix adjust | 75/243(31%) | 113/243(47%) | 28/243(12%) |
| | | | | | |
| --- | --- | --- | --- | --- | --- |
| Score | Expect | Method | Identities | Positives | Gaps |
| 67 bits(169) | 2e-12 | Compositional matrix adjust | 62/213(29%) | 108/213(51%) | 34/213(16%) |
e
Figure S1 supp: Protein BLAST from the human receptor sequences: Query: P36897 Length: 503 as a query against the respective Solanum lycopersicum species-specific protein sequence databases. Two proteins with significant homology to the human protein were identified: d. AOA3Q7F2Q7 (Solyc 02g072070), characterized as a serine-threonine kinase, and e. Q8GUQ5 (Solyc 04g051510), described as an LRR receptor kinase involved in tomato brassinosteroid (BR) hormone signaling.

### Slide 3
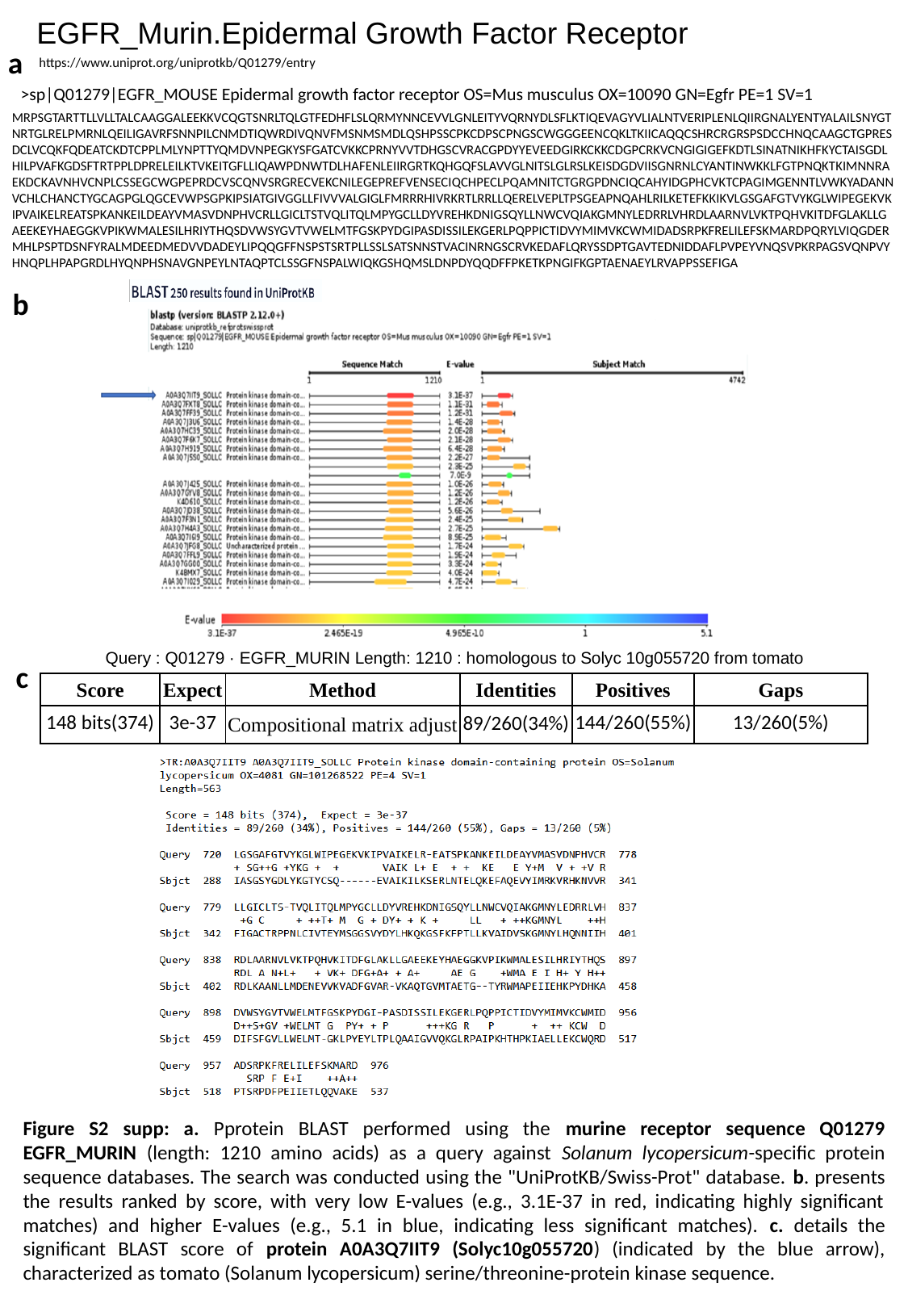

EGFR_Murin.Epidermal Growth Factor Receptor
a
https://www.uniprot.org/uniprotkb/Q01279/entry
>sp|Q01279|EGFR_MOUSE Epidermal growth factor receptor OS=Mus musculus OX=10090 GN=Egfr PE=1 SV=1
MRPSGTARTTLLVLLTALCAAGGALEEKKVCQGTSNRLTQLGTFEDHFLSLQRMYNNCEVVLGNLEITYVQRNYDLSFLKTIQEVAGYVLIALNTVERIPLENLQIIRGNALYENTYALAILSNYGTNRTGLRELPMRNLQEILIGAVRFSNNPILCNMDTIQWRDIVQNVFMSNMSMDLQSHPSSCPKCDPSCPNGSCWGGGEENCQKLTKIICAQQCSHRCRGRSPSDCCHNQCAAGCTGPRESDCLVCQKFQDEATCKDTCPPLMLYNPTTYQMDVNPEGKYSFGATCVKKCPRNYVVTDHGSCVRACGPDYYEVEEDGIRKCKKCDGPCRKVCNGIGIGEFKDTLSINATNIKHFKYCTAISGDLHILPVAFKGDSFTRTPPLDPRELEILKTVKEITGFLLIQAWPDNWTDLHAFENLEIIRGRTKQHGQFSLAVVGLNITSLGLRSLKEISDGDVIISGNRNLCYANTINWKKLFGTPNQKTKIMNNRAEKDCKAVNHVCNPLCSSEGCWGPEPRDCVSCQNVSRGRECVEKCNILEGEPREFVENSECIQCHPECLPQAMNITCTGRGPDNCIQCAHYIDGPHCVKTCPAGIMGENNTLVWKYADANNVCHLCHANCTYGCAGPGLQGCEVWPSGPKIPSIATGIVGGLLFIVVVALGIGLFMRRRHIVRKRTLRRLLQERELVEPLTPSGEAPNQAHLRILKETEFKKIKVLGSGAFGTVYKGLWIPEGEKVKIPVAIKELREATSPKANKEILDEAYVMASVDNPHVCRLLGICLTSTVQLITQLMPYGCLLDYVREHKDNIGSQYLLNWCVQIAKGMNYLEDRRLVHRDLAARNVLVKTPQHVKITDFGLAKLLGAEEKEYHAEGGKVPIKWMALESILHRIYTHQSDVWSYGVTVWELMTFGSKPYDGIPASDISSILEKGERLPQPPICTIDVYMIMVKCWMIDADSRPKFRELILEFSKMARDPQRYLVIQGDERMHLPSPTDSNFYRALMDEEDMEDVVDADEYLIPQQGFFNSPSTSRTPLLSSLSATSNNSTVACINRNGSCRVKEDAFLQRYSSDPTGAVTEDNIDDAFLPVPEYVNQSVPKRPAGSVQNPVYHNQPLHPAPGRDLHYQNPHSNAVGNPEYLNTAQPTCLSSGFNSPALWIQKGSHQMSLDNPDYQQDFFPKETKPNGIFKGPTAENAEYLRVAPPSSEFIGA
b
Query : Q01279 · EGFR_MURIN Length: 1210 : homologous to Solyc 10g055720 from tomato
| | | | | | |
| --- | --- | --- | --- | --- | --- |
| Score | Expect | Method | Identities | Positives | Gaps |
| 148 bits(374) | 3e-37 | Compositional matrix adjust | 89/260(34%) | 144/260(55%) | 13/260(5%) |
c
Figure S2 supp: a. Pprotein BLAST performed using the murine receptor sequence Q01279 EGFR_MURIN (length: 1210 amino acids) as a query against Solanum lycopersicum-specific protein sequence databases. The search was conducted using the "UniProtKB/Swiss-Prot" database. b. presents the results ranked by score, with very low E-values (e.g., 3.1E-37 in red, indicating highly significant matches) and higher E-values (e.g., 5.1 in blue, indicating less significant matches). c. details the significant BLAST score of protein A0A3Q7IIT9 (Solyc10g055720) (indicated by the blue arrow), characterized as tomato (Solanum lycopersicum) serine/threonine-protein kinase sequence.

### Slide 4
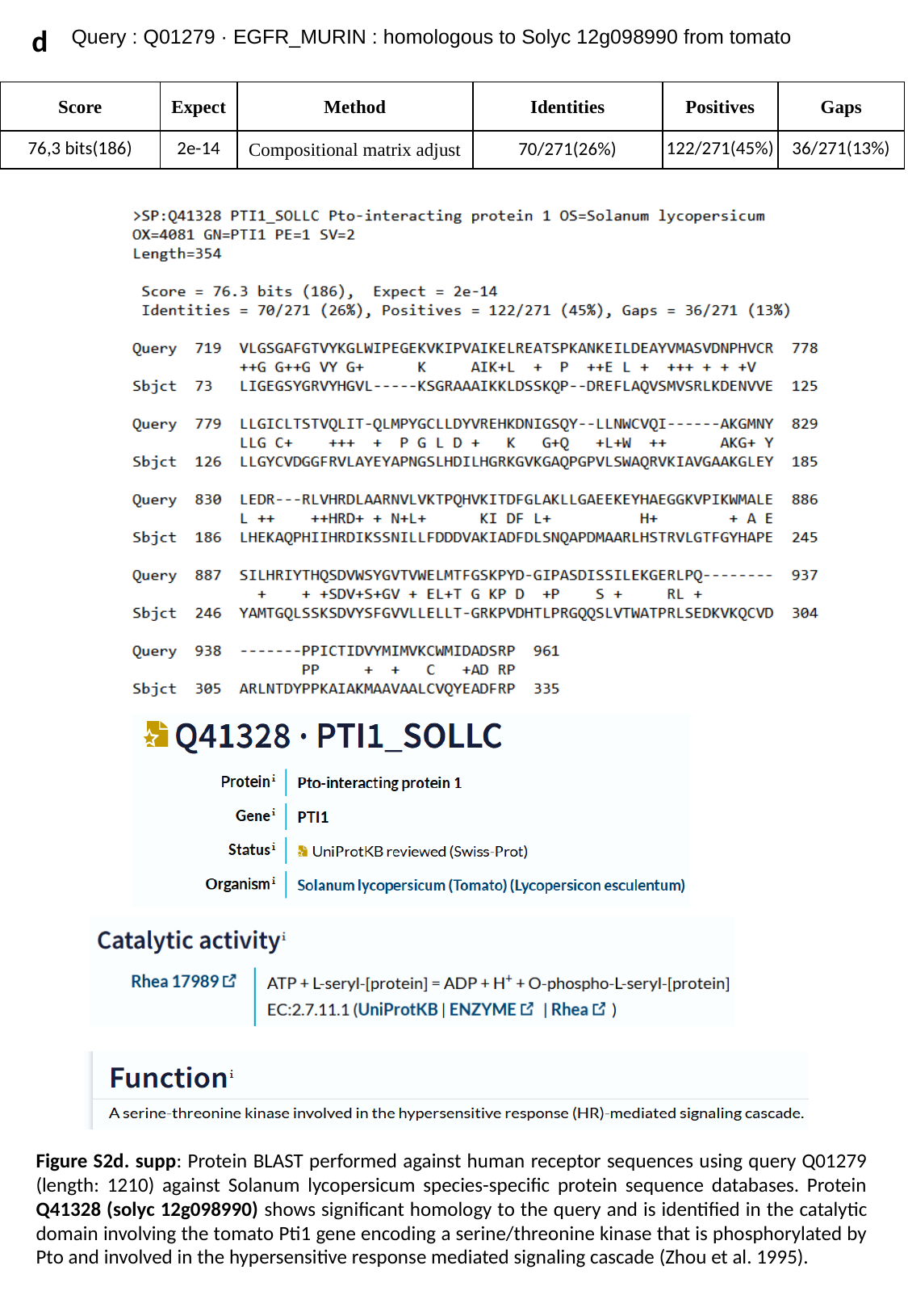

d
Query : Q01279 · EGFR_MURIN : homologous to Solyc 12g098990 from tomato
| | | | | | |
| --- | --- | --- | --- | --- | --- |
| Score | Expect | Method | Identities | Positives | Gaps |
| 76,3 bits(186) | 2e-14 | Compositional matrix adjust | 70/271(26%) | 122/271(45%) | 36/271(13%) |
Figure S2d. supp: Protein BLAST performed against human receptor sequences using query Q01279 (length: 1210) against Solanum lycopersicum species-specific protein sequence databases. Protein Q41328 (solyc 12g098990) shows significant homology to the query and is identified in the catalytic domain involving the tomato Pti1 gene encoding a serine/threonine kinase that is phosphorylated by Pto and involved in the hypersensitive response mediated signaling cascade (Zhou et al. 1995).

### Slide 5
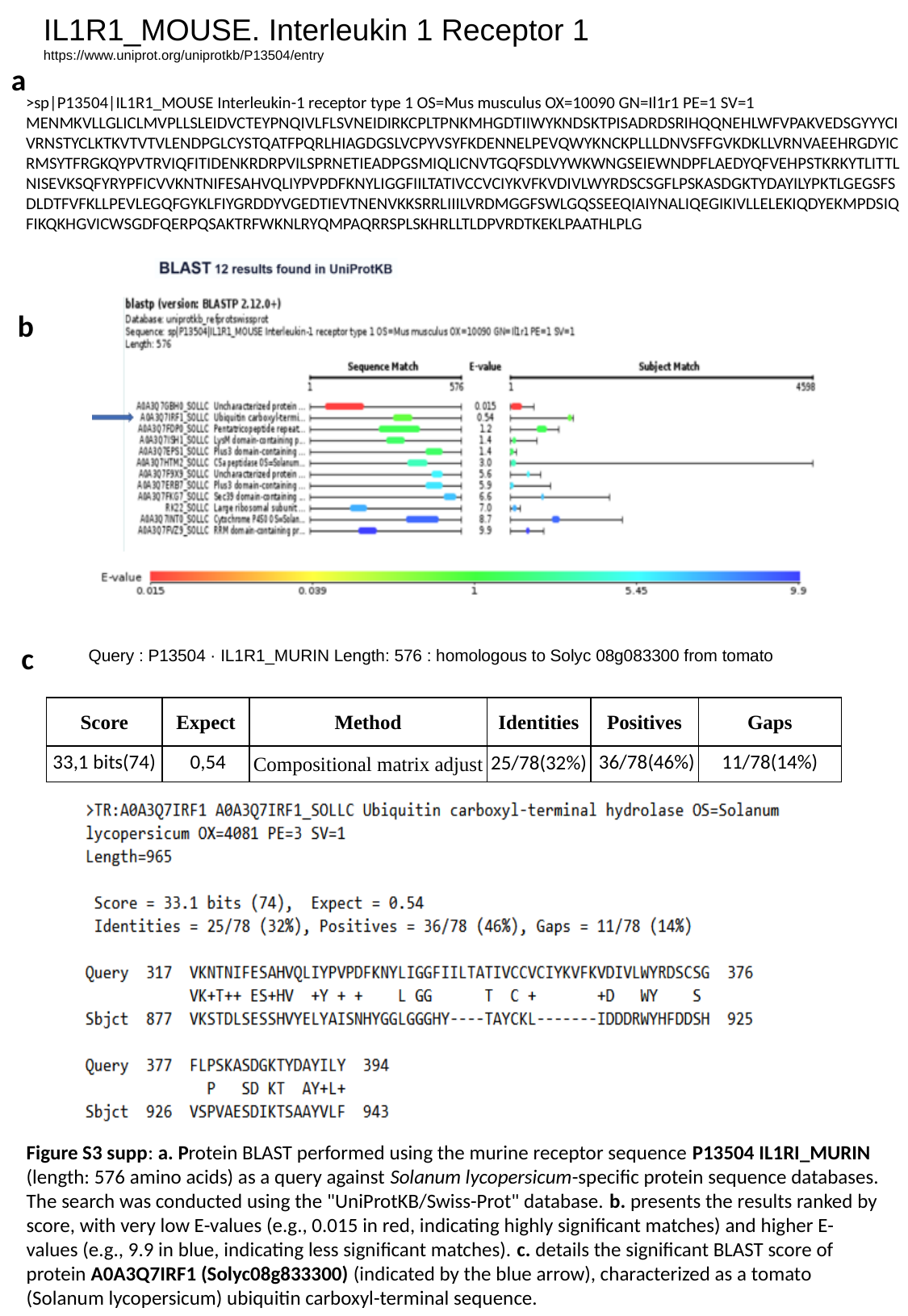

IL1R1_MOUSE. Interleukin 1 Receptor 1
https://www.uniprot.org/uniprotkb/P13504/entry
a
>sp|P13504|IL1R1_MOUSE Interleukin-1 receptor type 1 OS=Mus musculus OX=10090 GN=Il1r1 PE=1 SV=1
MENMKVLLGLICLMVPLLSLEIDVCTEYPNQIVLFLSVNEIDIRKCPLTPNKMHGDTIIWYKNDSKTPISADRDSRIHQQNEHLWFVPAKVEDSGYYYCIVRNSTYCLKTKVTVTVLENDPGLCYSTQATFPQRLHIAGDGSLVCPYVSYFKDENNELPEVQWYKNCKPLLLDNVSFFGVKDKLLVRNVAEEHRGDYICRMSYTFRGKQYPVTRVIQFITIDENKRDRPVILSPRNETIEADPGSMIQLICNVTGQFSDLVYWKWNGSEIEWNDPFLAEDYQFVEHPSTKRKYTLITTLNISEVKSQFYRYPFICVVKNTNIFESAHVQLIYPVPDFKNYLIGGFIILTATIVCCVCIYKVFKVDIVLWYRDSCSGFLPSKASDGKTYDAYILYPKTLGEGSFSDLDTFVFKLLPEVLEGQFGYKLFIYGRDDYVGEDTIEVTNENVKKSRRLIIILVRDMGGFSWLGQSSEEQIAIYNALIQEGIKIVLLELEKIQDYEKMPDSIQFIKQKHGVICWSGDFQERPQSAKTRFWKNLRYQMPAQRRSPLSKHRLLTLDPVRDTKEKLPAATHLPLG
b
c
Query : P13504 · IL1R1_MURIN Length: 576 : homologous to Solyc 08g083300 from tomato
| | | | | | |
| --- | --- | --- | --- | --- | --- |
| Score | Expect | Method | Identities | Positives | Gaps |
| 33,1 bits(74) | 0,54 | Compositional matrix adjust | 25/78(32%) | 36/78(46%) | 11/78(14%) |
Figure S3 supp: a. Protein BLAST performed using the murine receptor sequence P13504 IL1RI_MURIN (length: 576 amino acids) as a query against Solanum lycopersicum-specific protein sequence databases. The search was conducted using the "UniProtKB/Swiss-Prot" database. b. presents the results ranked by score, with very low E-values (e.g., 0.015 in red, indicating highly significant matches) and higher E-values (e.g., 9.9 in blue, indicating less significant matches). c. details the significant BLAST score of protein A0A3Q7IRF1 (Solyc08g833300) (indicated by the blue arrow), characterized as a tomato (Solanum lycopersicum) ubiquitin carboxyl-terminal sequence.

### Slide 6
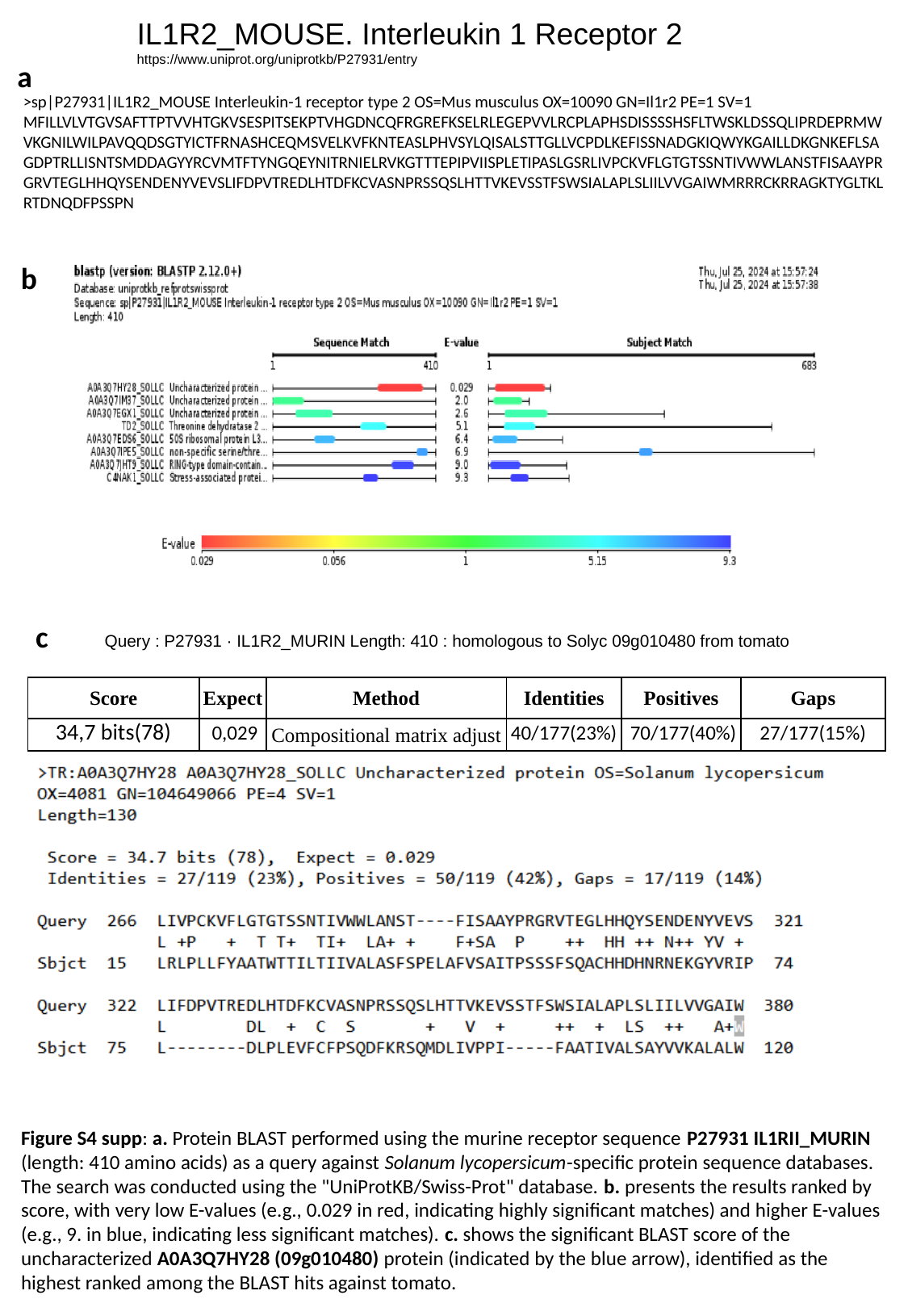

IL1R2_MOUSE. Interleukin 1 Receptor 2
https://www.uniprot.org/uniprotkb/P27931/entry
a
>sp|P27931|IL1R2_MOUSE Interleukin-1 receptor type 2 OS=Mus musculus OX=10090 GN=Il1r2 PE=1 SV=1
MFILLVLVTGVSAFTTPTVVHTGKVSESPITSEKPTVHGDNCQFRGREFKSELRLEGEPVVLRCPLAPHSDISSSSHSFLTWSKLDSSQLIPRDEPRMWVKGNILWILPAVQQDSGTYICTFRNASHCEQMSVELKVFKNTEASLPHVSYLQISALSTTGLLVCPDLKEFISSNADGKIQWYKGAILLDKGNKEFLSAGDPTRLLISNTSMDDAGYYRCVMTFTYNGQEYNITRNIELRVKGTTTEPIPVIISPLETIPASLGSRLIVPCKVFLGTGTSSNTIVWWLANSTFISAAYPRGRVTEGLHHQYSENDENYVEVSLIFDPVTREDLHTDFKCVASNPRSSQSLHTTVKEVSSTFSWSIALAPLSLIILVVGAIWMRRRCKRRAGKTYGLTKLRTDNQDFPSSPN
b
c
Query : P27931 · IL1R2_MURIN Length: 410 : homologous to Solyc 09g010480 from tomato
| | | | | | |
| --- | --- | --- | --- | --- | --- |
| Score | Expect | Method | Identities | Positives | Gaps |
| 34,7 bits(78) | 0,029 | Compositional matrix adjust | 40/177(23%) | 70/177(40%) | 27/177(15%) |
Figure S4 supp: a. Protein BLAST performed using the murine receptor sequence P27931 IL1RII_MURIN (length: 410 amino acids) as a query against Solanum lycopersicum-specific protein sequence databases. The search was conducted using the "UniProtKB/Swiss-Prot" database. b. presents the results ranked by score, with very low E-values (e.g., 0.029 in red, indicating highly significant matches) and higher E-values (e.g., 9. in blue, indicating less significant matches). c. shows the significant BLAST score of the uncharacterized A0A3Q7HY28 (09g010480) protein (indicated by the blue arrow), identified as the highest ranked among the BLAST hits against tomato.

### Slide 7
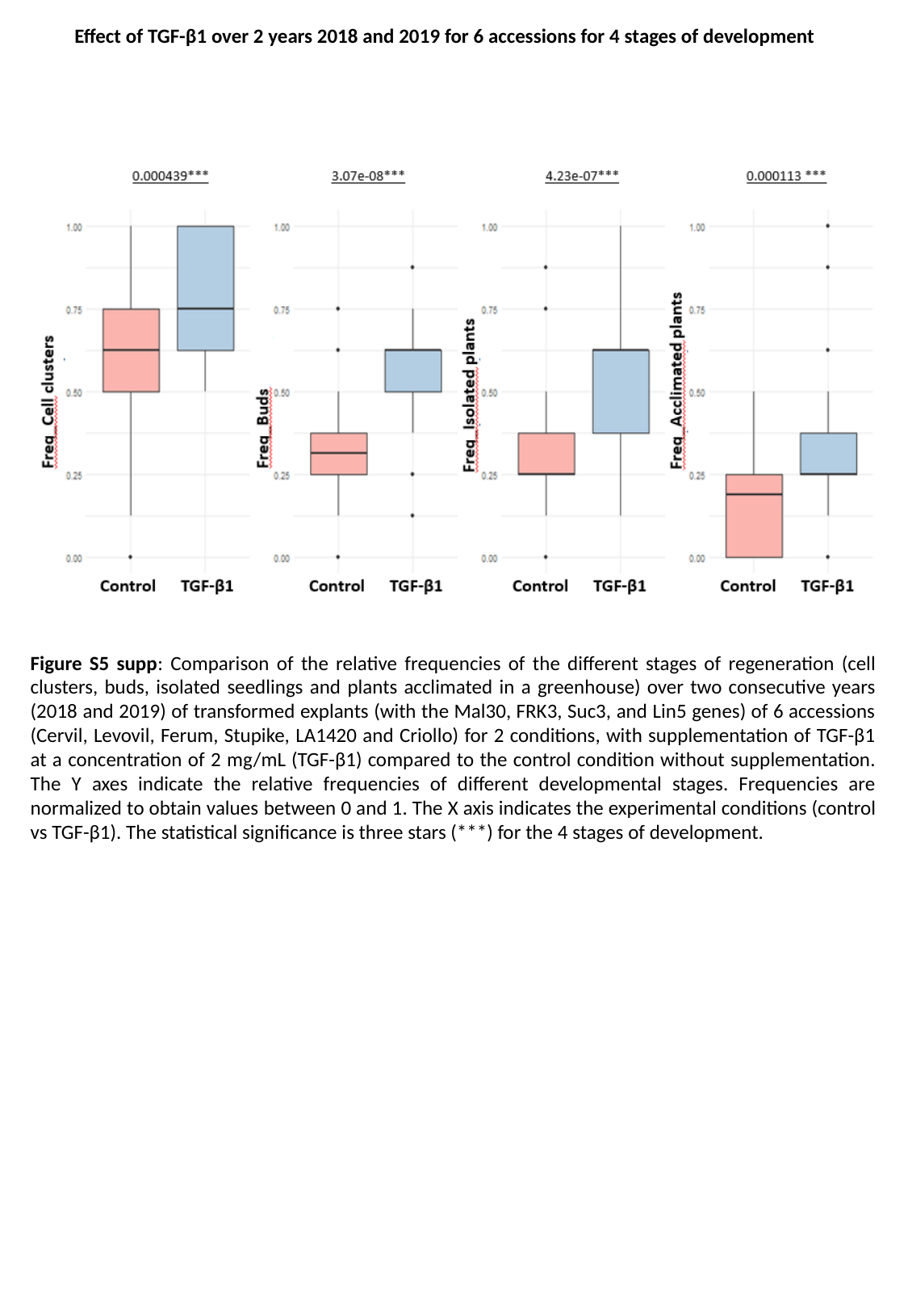

Effect of TGF-β1 over 2 years 2018 and 2019 for 6 accessions for 4 stages of development
Figure S5 supp: Comparison of the relative frequencies of the different stages of regeneration (cell clusters, buds, isolated seedlings and plants acclimated in a greenhouse) over two consecutive years (2018 and 2019) of transformed explants (with the Mal30, FRK3, Suc3, and Lin5 genes) of 6 accessions (Cervil, Levovil, Ferum, Stupike, LA1420 and Criollo) for 2 conditions, with supplementation of TGF-β1 at a concentration of 2 mg/mL (TGF-β1) compared to the control condition without supplementation. The Y axes indicate the relative frequencies of different developmental stages. Frequencies are normalized to obtain values ​​between 0 and 1. The X axis indicates the experimental conditions (control vs TGF-β1). The statistical significance is three stars (***) for the 4 stages of development.

### Slide 8
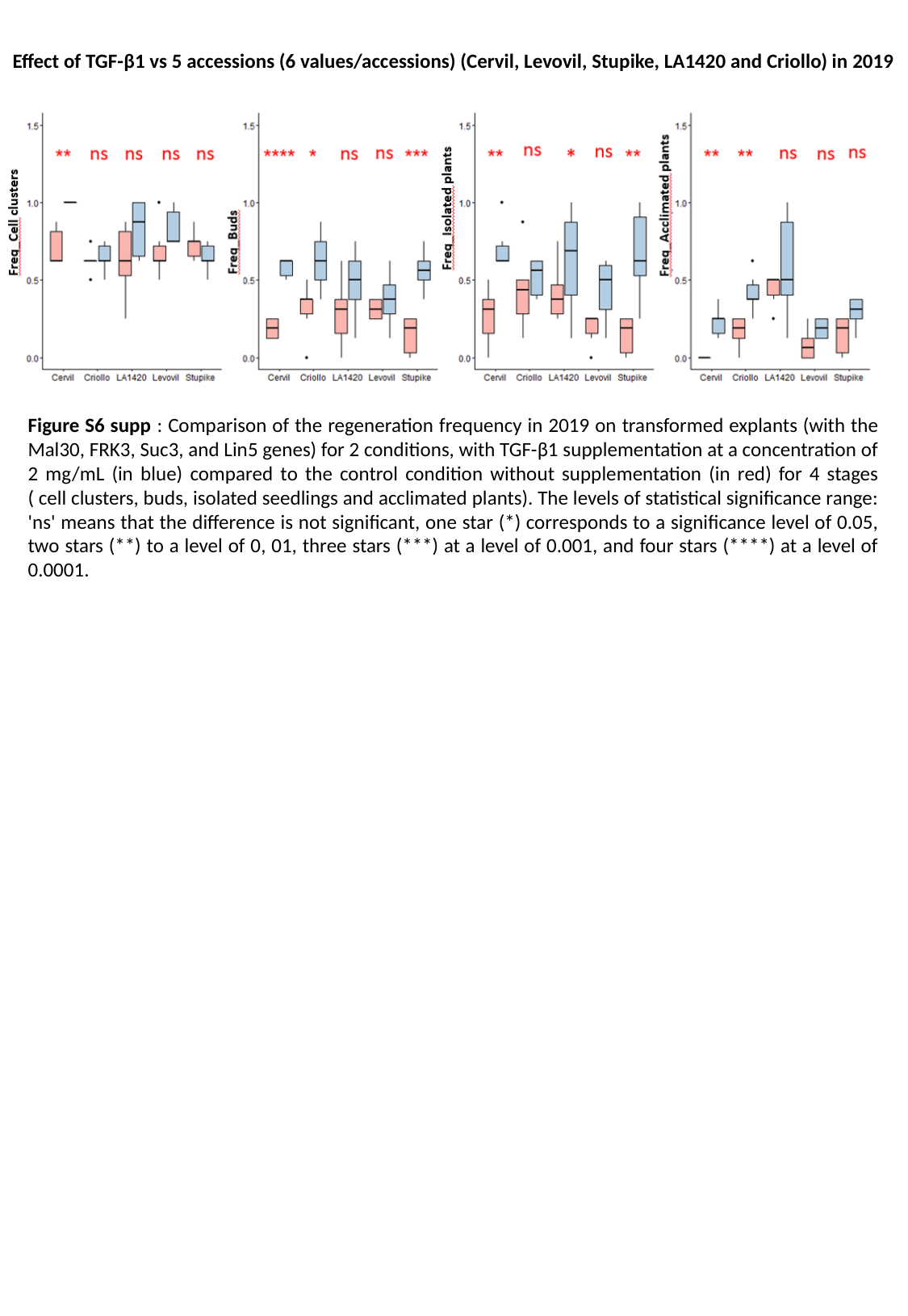

Effect of TGF-β1 vs 5 accessions (6 values/accessions) (Cervil, Levovil, Stupike, LA1420 and Criollo) in 2019
Figure S6 supp : Comparison of the regeneration frequency in 2019 on transformed explants (with the Mal30, FRK3, Suc3, and Lin5 genes) for 2 conditions, with TGF-β1 supplementation at a concentration of 2 mg/mL (in blue) compared to the control condition without supplementation (in red) for 4 stages ( cell clusters, buds, isolated seedlings and acclimated plants). The levels of statistical significance range: 'ns' means that the difference is not significant, one star (*) corresponds to a significance level of 0.05, two stars (**) to a level of 0, 01, three stars (***) at a level of 0.001, and four stars (****) at a level of 0.0001.

### Slide 9
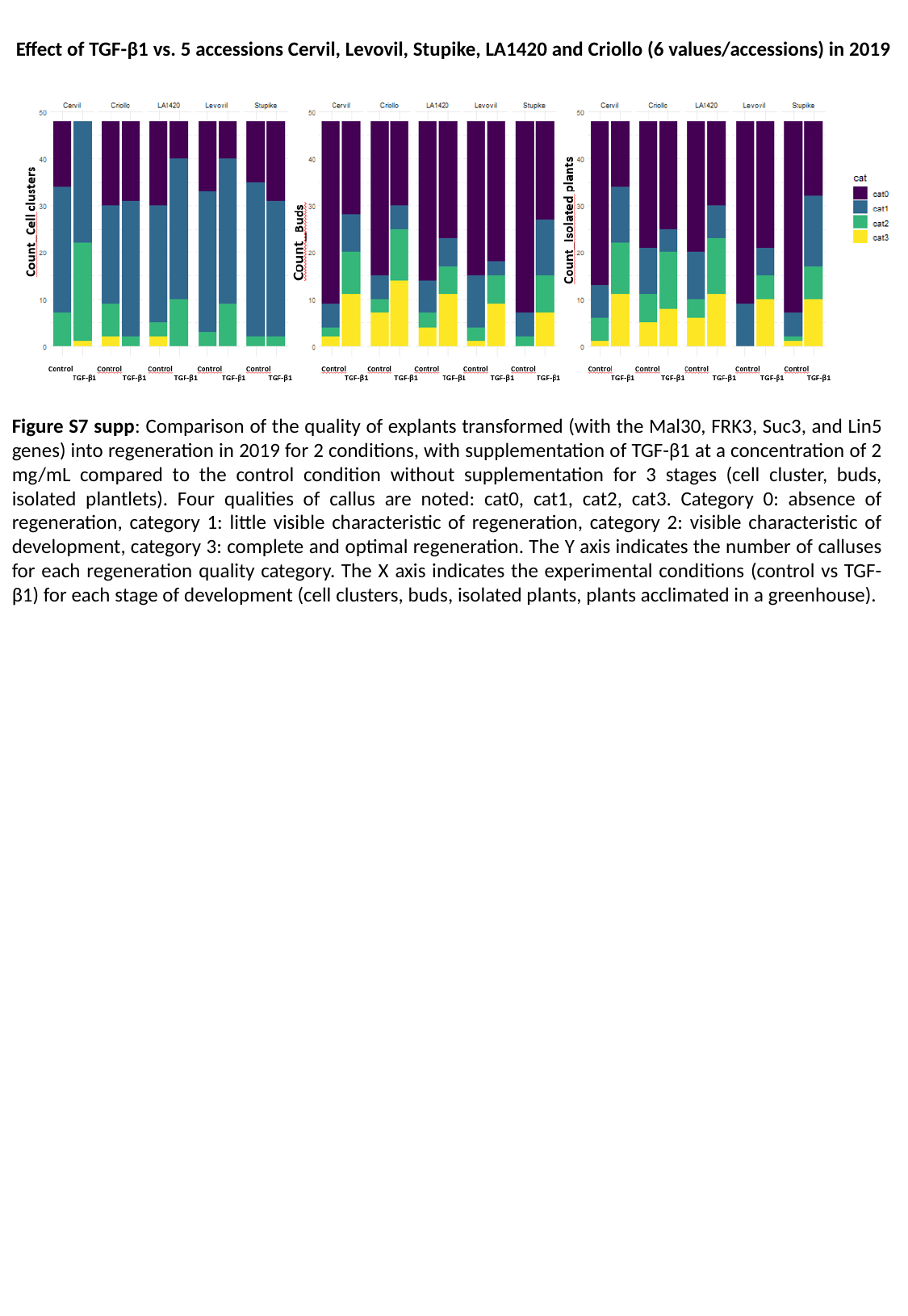

Effect of TGF-β1 vs. 5 accessions Cervil, Levovil, Stupike, LA1420 and Criollo (6 values/accessions) in 2019
Figure S7 supp: Comparison of the quality of explants transformed (with the Mal30, FRK3, Suc3, and Lin5 genes) into regeneration in 2019 for 2 conditions, with supplementation of TGF-β1 at a concentration of 2 mg/mL compared to the control condition without supplementation for 3 stages (cell cluster, buds, isolated plantlets). Four qualities of callus are noted: cat0, cat1, cat2, cat3. Category 0: absence of regeneration, category 1: little visible characteristic of regeneration, category 2: visible characteristic of development, category 3: complete and optimal regeneration. The Y axis indicates the number of calluses for each regeneration quality category. The X axis indicates the experimental conditions (control vs TGF-β1) for each stage of development (cell clusters, buds, isolated plants, plants acclimated in a greenhouse).

### Slide 10
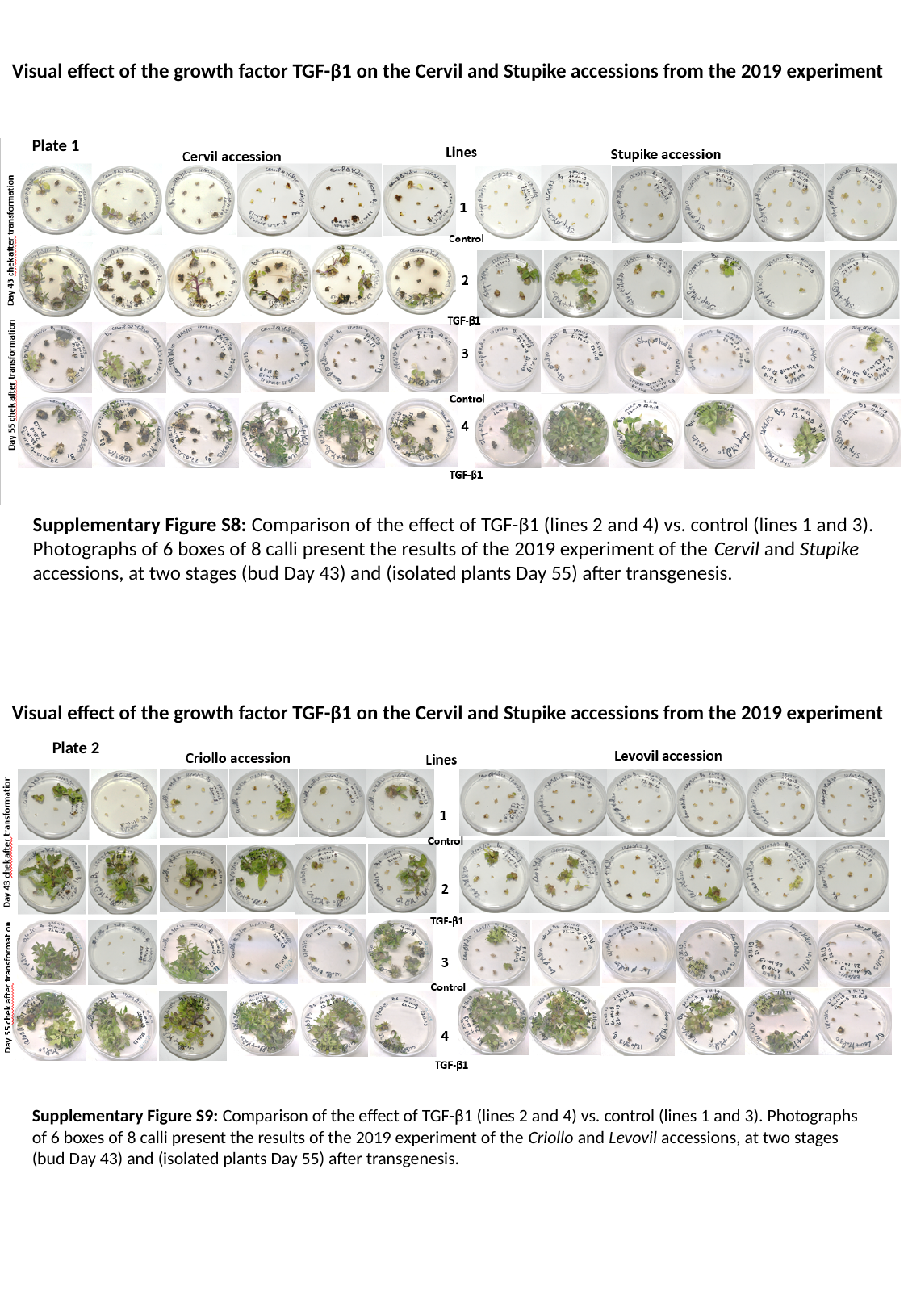

Visual effect of the growth factor TGF-β1 on the Cervil and Stupike accessions from the 2019 experiment
Plate 1
Supplementary Figure S8: Comparison of the effect of TGF-β1 (lines 2 and 4) vs. control (lines 1 and 3). Photographs of 6 boxes of 8 calli present the results of the 2019 experiment of the Cervil and Stupike accessions, at two stages (bud Day 43) and (isolated plants Day 55) after transgenesis.
Visual effect of the growth factor TGF-β1 on the Cervil and Stupike accessions from the 2019 experiment
Plate 2
Supplementary Figure S9: Comparison of the effect of TGF-β1 (lines 2 and 4) vs. control (lines 1 and 3). Photographs of 6 boxes of 8 calli present the results of the 2019 experiment of the Criollo and Levovil accessions, at two stages (bud Day 43) and (isolated plants Day 55) after transgenesis.

### Slide 11
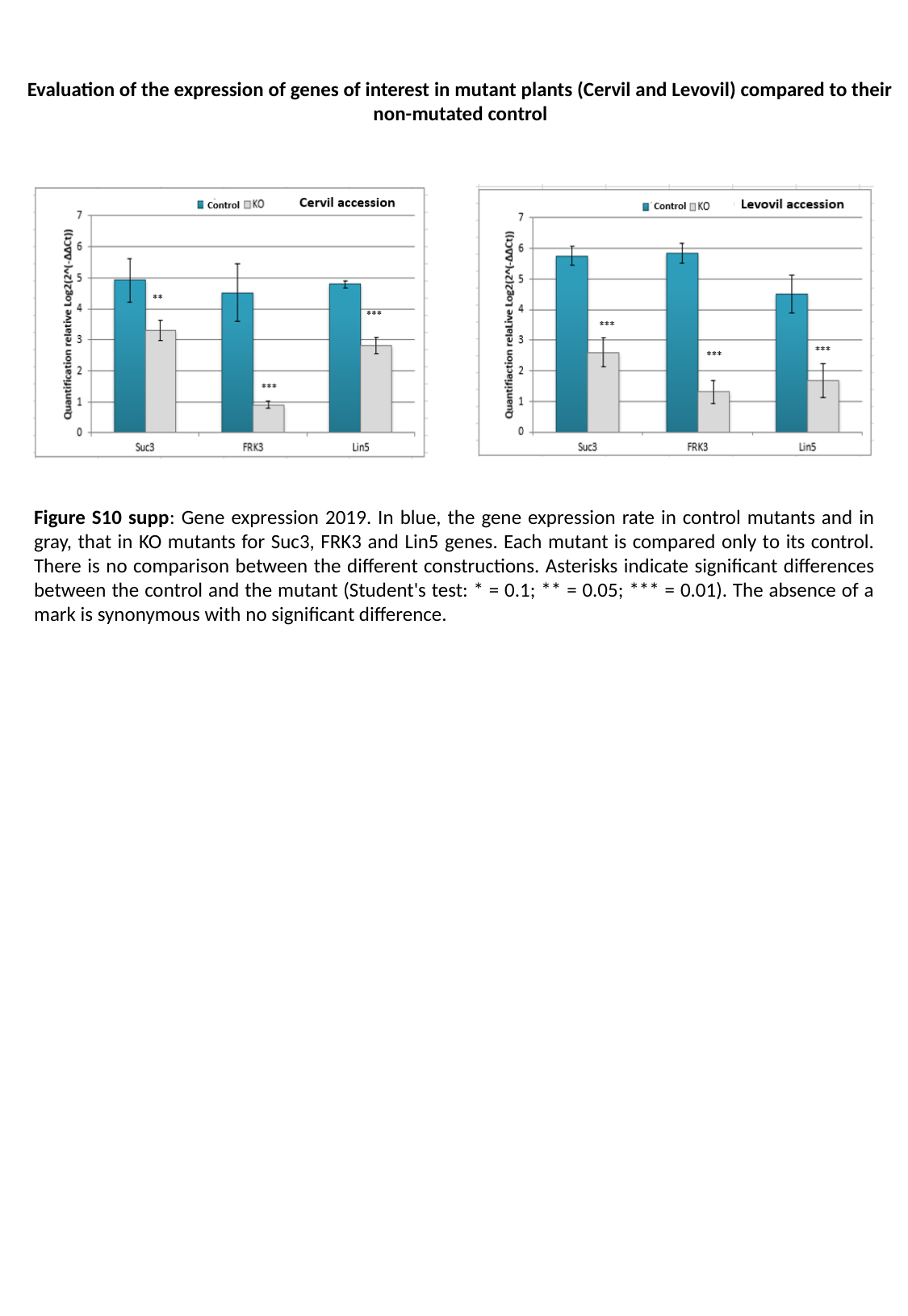

Evaluation of the expression of genes of interest in mutant plants (Cervil and Levovil) compared to their non-mutated control
Figure S10 supp: Gene expression 2019. In blue, the gene expression rate in control mutants and in gray, that in KO mutants for Suc3, FRK3 and Lin5 genes. Each mutant is compared only to its control. There is no comparison between the different constructions. Asterisks indicate significant differences between the control and the mutant (Student's test: * = 0.1; ** = 0.05; *** = 0.01). The absence of a mark is synonymous with no significant difference.

### Slide 12
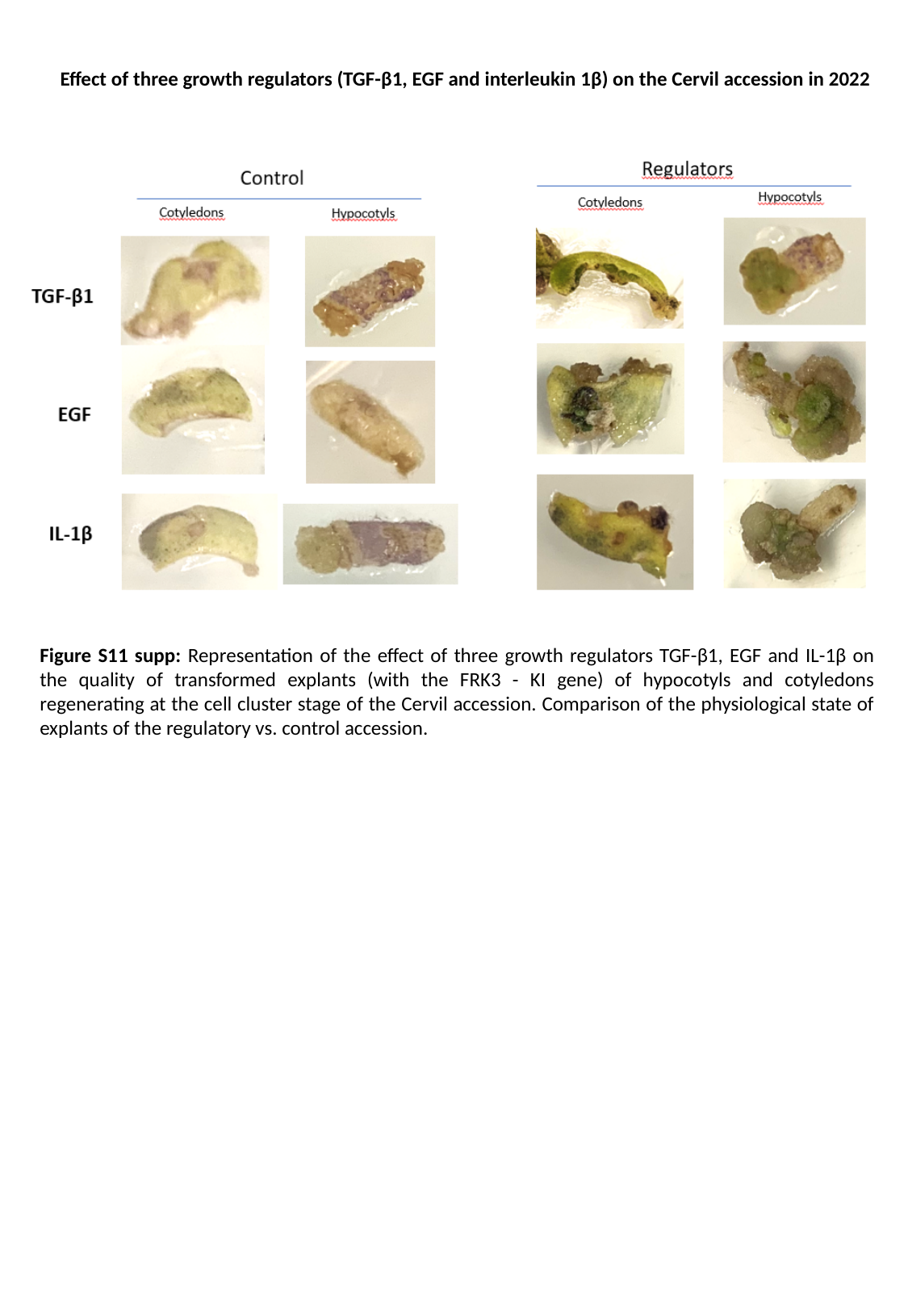

Effect of three growth regulators (TGF-β1, EGF and interleukin 1β) on the Cervil accession in 2022
Figure S11 supp: Representation of the effect of three growth regulators TGF-β1, EGF and IL-1β on the quality of transformed explants (with the FRK3 - KI gene) of hypocotyls and cotyledons regenerating at the cell cluster stage of the Cervil accession. Comparison of the physiological state of explants of the regulatory vs. control accession.
