## supplementary tables for "Mammalian growth-regulating factors enhance regeneration of recalcitrant transgenic tomato accessions"

**Table 1 : details of constructs**

**a - Knock Out Constructs:**

| **Sequence gRNAs Malate Solyc 06g072930: 1832 bp** | | | | | | | |
| --- | --- | --- | --- | --- | --- | --- | --- |
| **gRNA 1 5’CTTCCCAACTAGCTTCTTGCTTTCTCCTCGCAGCACGGGACACTTTACTTATTTGCTATCATCACTACAAATTAAACCTAAAAGTAGCAATTTCAACTTGCTCAACACTTGCCCTT3’** | | | | | | | |
| **gRNA 2**  **5’GCACCTTACTTCCCCATTGGACATGTGCTATTTTAATTGAGCACTTTTGTATTTTCATTTCTCCTTTGCTCAAAAAGCACTCTTCATAACGGCAGATTATAATCGCAATGTGGACGGGAAATAATGTGACCAGGCCATTTAGGGTTGCAGGAGCAGCTGCT3’** | | | | | | | |
| **Features :** | | | | | | | |
| **gRNA1 KO Mal30 : [89 : 111]**  **gRNA2 KO Mal30 : [1275 : 1297]** | | **Primer Mal30F gRNA1 : [64 : 83]**  **Primer Mal30KO1R gRNA1 : [160 : 179]** | | | **Primer Mal30_KO2_F gRNA2 : [1180 : 1199]**  **Primer Mal30R gRNA2 : [1310 : 1329]** | | |
| **Sequence gRNAs Invertase 03 Solyc 03g083910 : 5477 bp** | | | | | | | |
| **gRNA 1 - gRNA 2**  **5’CTTGCGTACCCCGCCAACTTATCTGATCCTCTCCTTCTAGACTGGGTCAAGTTCAAAGGCAACCCGGTTCTGGTTCCTCCACCCGGCATTGGTGTCAAGGACTTTAGAGACCCGACTACTGCTTGGACCGGACCACAAAATGGGCAATGGCTGTTAACAATCGGGTCTAAGATTGGTAAAACGGGTGTTGCACTTGTTTATGAAACTTCCAACTTCACAAGCTTTAAGCTATTGGATGGAGTGCTGCATGCGGTTCCGGGTACGGGTATGTGGGAGTGTGTGGACTTTTACCCGGTATCTACTAAAAAAA3’** | | | | | | | |
| **Features :** | | | | | | | |
| **gRNA 1 KO Inv03g: [2922: 2940]**  **gRNA 2 KO Inv03g: [2951 : 2973]** | | | | **Primer F Inv03: [2850 : 2870]**  **Primer R Inv03: [3105 : 3124]** | | | |
| **Sequence gRNAs Invertase 09 Solyc 09g010080 : 5064 bp** | | | | | | | |
| **gRNA 1**  **5’CAAACCTGTTATCATATACACCGGAGTAGTAGATTCGTATAATAATCAAGTCCAGAACTACGCCATCCCGGCTAACCTATCTGATCCATTTCTTCGTAAATGGATCAAACCTAACAACAACCCGTTGATCGTCCCTGATAACAGTATCAA3’** | | | | | | | |
| **gRNA 2**  **5’TCAAAGCCCAACATCCACTTCATTCATCTACTAATACTGGAAATTGGGAGTGTCCTGATTTTTTCCCTGTATTATTTAATAGTACCAATGGTTTAGATGTATCGTATCGCGGAAAAAATGTTAAATATGTCCTCAAGAATAGTCTTGATGTTGCTAGGTTTGATTATTACACTATTGGCATGTATCACACCAAAAT3’** | | | | | | | |
| **Features :** | | | | | | | |
| **gRNA1 KO Inv09g: [2460 : 2482]**  **gRNA2 KO Inv09g: [2782 : 2804]** | | | **Primer Inv09_F: [2429 : 2449]**  **Primer Inv09_R: [2516 : 2536]** | | | **Primer Inv09_F: [2699 : 2721]**  **Primer Inv09_R : [2833 : 2855]** | |
| **Sequence gRNAs Invertase Fructokinase 3 (FRK3) Solyc 02g091490 : 5950 bp** | | | | | | | |
| **gRNA 1**  **5’ACTTTCTAATAAATGGCTCTTCATGCTACTGCTTTCTCCTTCACTGGAGTTTCTACTTCAAGTAAATCTTCCAGAAGCGCTTTGCTTTCAGTCTTTCCTCTTCCTAGAAGATGCACTGTCAAAGCAACTTCCCAGTATCCGCACAGCTTTCCTCGATGTAAAATCCAAGGTGTTCATCTTTTTAAATTTAGTCTACTTTC3’** | | | | | | | |
| **gRNA 2**  **5’TTGCCAAGTGACAATGGGCTAGTGGAGAAGGATGAATCTTCTCTTGTTGTGTGCTTTGGAGAAATGCTCATTGATTTTGTTCCGACTACAAGTGGGCTTTCATTGGCTGAAGCTCCTGCATTTAAAAAGGCTCCTGGTGGTGCACCAGCTAATGTTGCTGTTGGTATTTCCCGTCTTGGTGGTTCATCAGCTTTCATTGG3’** | | | | | | | |
| **Features :** | | | | | | | |
| **gRNA 1 KO FRK3: [1032: 1054]**  **gRNA 2 KO FRK3: [1282 : 1304]** | **Primer FRK3 F: [914:928]**  **Primer FRK3 R: [1054:1071]** | | | | | | **Primer FRK3 F : [1215 : 1229]**  **Primer FRK3 R : [1360 : 1374]** |

**b – Base editing Construct:**

**Sequence gRNAs Invertase Fructokinase 3 (FRK3) Solyc 02g091490 : 5950 bp**

**FRK3-KI-1: sL2.5-52770977 (c/a)**

ggggacaagtttgtacaaaaaagcaggcttc**CCATTTATATGGGAAAGAACAATAGTATTTCTTATATAGGCCCATTTAAGTTGAAAACAATCTTCAAAAGTCCCACATCGCTTAGATAAGAAAACGAAGCTGAGTTTATATACAGCTAGAGTCGAAGTAGTGATTGT**TAGAAAGTAAAAAAACCAG**GTTTTAGAGCTAGAAATAGCAAGTTAAAATAAGGCTAGTCCGTTATCAACTTGAAAAAGTGGCACCGAGTCGGTGC**TTTTTTTCTATC**ACTAGT**gacccagctttcttgtacaaagtggtcccc

**FRK3-KI-3: sL2.5-52766683(c/t)**

ggggacaagtttgtacaaaaaagcaggcttc**CCATTTATATGGGAAAGAACAATAGTATTTCTTATATAGGCCCATTTAAGTTGAAAACAATCTTCAAAAGTCCCACATCGCTTAGATAAGAAAACGAAGCTGAGTTTATATACAGCTAGAGTCGAAGTAGTGATTG**AGAACTATGCAAGGAGCACA**GTTTTAGAGCTAGAAATAGCAAGTTAAAATAAGGCTAGTCCGTTATCAACTTGAAAAAGTGGCACCGAGTCGGTGC**TTTTTTTCTATC**ACTAGT**gacccagctttcttgtacaaagtggtcccc

**Table 2 : Map of plasmids used for construct cloning**

**a - Complete sequence map of the binary expression plasmid pDe Cas9 used for the knockout constructs**

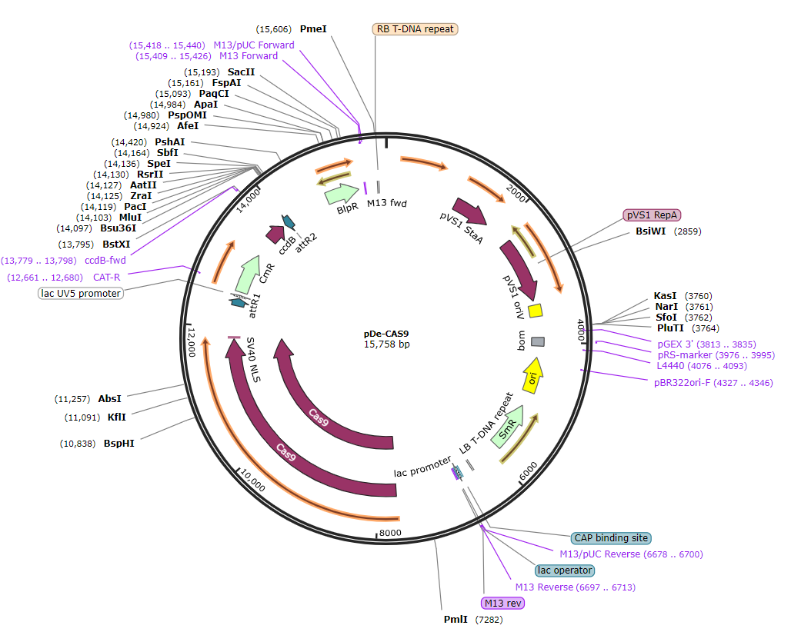

**b - Complete sequence map of the binary expression plasmid pDicAID_nCas9-PmCDA_NptII_Della used for the Knock in constructs**

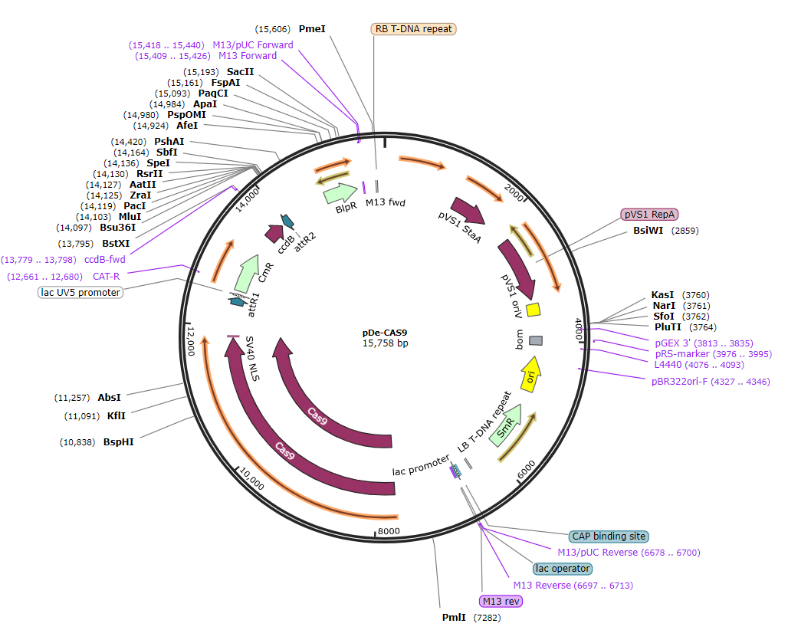

**Table 3 : Protein sequences of the regulators**

| **Recombinant human transforming growth factor β1** | [ALDTNYCFSS TEKNCCVRQL YIDFRKDLGW KWIHEPKGYH ANFCCLGPCPY IWSLDTQYSK VLALYNQHNP GASAAPCCVP QALEPLPIVY YVGRKPKVEQ LSNMIVRSCK CS] |
| --- | --- |
| **Recombinant murine epidermal growth factor EGF** | [NSYPGCPSSY DGYCLNGGVC MHIESLDSYT CNCVIGYSGD RCQTRDLRWW ELR] |
| **Recombinant murine interleukin IL-1β** | [MVPIRQLHYR LRDEQQKSLV LSDPYELKAL HLNGQNINQQ VIFSMSFVQG EPSNDKIPVA LGLKGKNLYL SCVMKDGTPT LQLESVDPKQ YPKKKMEKRF VFNKIEVKSK VEFESAEFPN WYISTSQAEH KPVFLGNNSG QDIIDFTMES] |

**Table 4 : Composition of media used for the culture of agrobacteria and the in vitro culture of calluses and transgenic plants**

| **Lysogeny selective broth medium** | **Coculture medium** |
| --- | --- |
| **Composition for 1 liter** | |
| NaCl 10g | Macroelements (x20) 50 mL |
| Bactotrytone 20g | Microelements (x1000) 1mL |
| Yeast extract 5g | Nitsch Vitamins (x100) 10mL |
| Agar (for solid medium only) 12g | KH₂PO₄ 200mg |
| q.s. distilled water | Iron EDTA (x200) 5mL |
|  | Sucrose 30g |
| Adjusted the PH to 7.5  Added with rifampicin (25mg/l), ampicillin (50mg/l) selection of entry vector, spectinomycin (100mg/l) selection of expression vector for Agrobacterium tumefaciens strain 'C58' and the helper plasmid 'pGV2260' | Adjusted the PH to 5.6 - 6  Added with vitamin B1 (0.9mg/l), 2,4D (0.2mg/l), kinetin (0.1mg/l), acetosyringone (0.2mM), growth regulators (efficacy from 0.1 of 20 ng/mL) |
| **Selective Murashige and Skoog medium** | **Selective Murashige and Skoog ½ medium** |
| **Composition for 1 liter** | |
| Macroelements (x20) 50mL | Macroelements (x20) 25mL |
| Microelements (x1000) 1mL | Microelements (x1000) 0.5mL |
| Nitsch Vitamins (x100) 10mL | Nitsch Vitamins (x100) 10mL |
| Iron EDTA (x200) 5mL | Iron EDTA (x200) 5mL |
| Sucrose 30g | Sucrose 30g |
| Agar 7g | Agar 7g |
| Adjusted the PH to 5.6 - 6  Added with kanamycin (100mg/l), ticarcillin (225mg/l), zeatin (2mg/l) | Adjusted the PH to 5.6 - 6  Added with kanamycin (100mg/l), ticarcillin (225mg/l) |

**Table 5 : Genome editing results of secondary transformants of Cervil and Levovil accessions obtained with the addition of TGF during transgenesis for the target genes Suc3, FRK3 and Lin5** (- no muted).

| **Name** | **Mutation on gRNA1** | **Mutation on gRNA2** |
| --- | --- | --- |
| T Cervil Suc3 |  |  |
| Suc3-Cer1.1 | - | - |
| Suc3-Cer1.2 | Insertion of a T | Insertion of a T |
| Suc3-Cer1.3 | - | - |
| Suc3-Cer1.4 | - | - |
| Suc3-Cer1.5 | - | - |
| Suc3-Cer1.6 | - | - |
| Suc3-Cer1.7 | - | - |
| Suc3-Cer1.8 | - | - |
| Suc3-Cer1.9 | - | - |
| Suc3-Cer1.10 | - | - |
| Suc3-Cer1.11 | - | - |
| Suc3-Cer1.12 | - | - |
| Suc3-Cer1.13 | Insertion of a T | Insertion of a T |
| Suc3-Cer1.14 | Insertion of a T | Insertion of a T |
| Suc3-Cer1.15 | Insertion of a T | Insertion of a T |
| Suc3-Cer1.16 | Insertion of a T | Insertion of a T |
| Suc3-Cer1.17 | - | - |
| Suc3-Cer1.18 | Insertion of a T | Insertion of a T |
| Suc3-Cer1.19 | Insertion of a T | Insertion of a T |
| Suc3-Cer1.20 | Insertion of a T | Insertion of a T |
| Suc3-Cer1.21 | Insertion of a T | Insertion of a T |
| Suc3-Cer1.22 | Insertion of a T | Insertion of a T |
| Suc3-Cer1.23 | Insertion of a T | Insertion of a T |
| Suc3-Cer1.24 | Insertion of a T | Insertion of a T |
| Suc3-Cer1.25 | - | - |
| Suc3-Cer2.1 | - | - |
| Suc3-Cer2.2 | No alignment | Deletion of an A |
| Suc3-Cer2.3 | - | - |
| Suc3-Cer2.4 | - | - |
| Suc3-Cer2.5 | - | - |
| Suc3-Cer2.6 | No alignment | Deletion of an A |
| Suc3-Cer2.7 | - | - |
| Suc3-Cer2.8 | - | - |
| Suc3-Cer2.10 | - | - |
| Suc3-Cer2.11 | No alignment | Deletion of an A |
| Suc3-Cer2.12 | No alignment | Deletion of an A |
| Suc3-Cer2.13 | No alignment | Deletion of an A |
| Suc3-Cer2.14 | - | - |
| Suc3-Cer2.15 | No alignment | Deletion of an A |
| Suc3-Cer2.16 | No alignment | Deletion of an A |
| Suc3-Cer2.17 | No alignment | Deletion of an A |
| Suc3-Cer2.18 | No alignment | Deletion of an A |
| Suc3-Cer2.19 | No alignment | Deletion of an A |
| Suc3-Cer2.20 | No alignment | Deletion of an A |
| Suc3-Cer2.21 | No alignment | Deletion of an a |
| Suc3-Cer2.22 | No alignment | Deletion of an a |
| Suc3-Cer2.23 | No alignment | Deletion of an a |
| Suc3-Cer3.1 | No alignment | Deletion of an a |
| Suc3-Cer3.2 | No alignment | Deletion of an a |
| Suc3-Cer3.3 | - | - |
| Suc3-Cer3.4 | - | - |
| Suc3-Cer3.5 | - | - |
| Suc3-Cer3.6 | No alignment | Deletion of an a |
| Suc3-Cer3.7 | No alignment | Deletion of an a |
| Suc3-Cer3.8 | No alignment | Deletion of an a |
| Suc3-Cer3.9 | No alignment | Deletion of an a |
| Suc3-Cer3.10 | No alignment | Deletion of an a |
| Suc3-Cer3.11 | - | - |
| Suc3-Cer3.12 | - | - |
| Suc3-Cer3.13 | - | - |
| Suc3-Cer3.16 | - | - |
| Suc3-Cer3.17 | - | - |
| Suc3-Cer3.18 | - | - |
| Suc3-Cer3.19 | - | - |
| Suc3-Cer3.20 | - | - |
| Suc3-Cer3.21 | - | - |
| Suc3-Cer3.22 | - | - |
| Suc3-Cer3.23 | - | - |

| **Name** | **Mutation on gRNA1** | **Mutation on gRNA2** |
| --- | --- | --- |
| T Cervil FRK3 |  |  |
| FRK3-Cer2.1 | - | - |
| FRK3-Cer2.2 | Deletion of an A | Insertion of a T |
| FRK3-Cer2.3 | several indefinite bases | Insertion of a T and an A |
| FRK3-Cer2.4 | Inserting a C | Inserting an A |
| FRK3-Cer2.5 | several indefinite bases | Insertion of a T and an A |
| FRK3-Cer2.6 | Inserting a C | Inserting an A |
| FRK3-Cer2.7 | several indefinite bases | Insertion of a T and an A |
| FRK3-Cer2.8 | Inserting a C | Inserting an A |
| FRK3-Cer2.9 | Inserting a C | Inserting an A |
| FRK3-Cer2.10 | several indefinite bases | Insertion of a T and an A |
| FRK3-Cer2.11 | - | - |
| FRK3-Cer2.13 | Inserting a C | Insertion of an A and a G |
| FRK3-Cer2.14 | Inserting a C | Insertion of an A |
| FRK3-Cer2.15 | several indefinite bases | Insertion of a T and an A |
| FRK3-Cer2.16 | - | - |
| FRK3-Cer2.17 | - | - |
| FRK3-Cer2.18 | - | - |
| FRK3-Cer2.19 | - | - |
| FRK3-Cer2.20 | several indefinite bases | Insertion of a T and an A |
| FRK3-Cer2.21 | - | - |
| FRK3-Cer2.22 | several indefinite bases | Insertion of a T and an A |
| FRK3-Cer2.23 | several indefinite bases | Insertion of a T and an A |
| FRK3-Cer2.24 | Deletion of an A | Insertion of a T |
| FRK3-Cer2.25 | Deletion of an A | Insertion of a T |
| FRK3-Cer3.1 | - | - |
| FRK3-Cer3.2 | Insertion of a T | Insertion of an A |
| FRK3-Cer3.3 | Insertion of a T | Insertion of an A |
| FRK3-Cer3.4 | Insertion of a T | Insertion of an A |
| FRK3-Cer3.5 | Insertion of a T | Insertion of an A |
| FRK3-Cer3.6 | - | - |
| FRK3-Cer3.7 | - | - |
| FRK3-Cer3.8 | - | - |
| FRK3-Cer3.9 | Insertion of a T | Insertion of an A |
| FRK3-Cer3.10 | - | - |
| FRK3-Cer3.11 | - | - |
| FRK3-Cer3.12 | - | - |
| FRK3-Cer3.13 | - | - |
| FRK3-Cer3.14 | - | - |
| FRK3-Cer3.15 | - | - |
| FRK3-Cer3.16 | - | - |
| FRK3-Cer3.17 |  | - |
| FRK3-Cer3.18 | Insertion of a T | Insertion of an A |
| FRK3-Cer3.19 | - | - |
| FRK3-Cer3.20 | - | - |
| FRK3-Cer3.21 | - | - |
| FRK3-Cer3.22 | - | - |
| FRK3-Cer3.23 | Insertion of a T | Insertion of an A |
| FRK3-Cer3.24 | - | - |

| **Name** | **Mutation on gRNA1** | **Mutation on gRNA2** |
| --- | --- | --- |
| T Cervil Lin5 |  |  |
| Lin5-Cer3.1 | Deletion of a T and insertion of a GCG sequence | Deletion of an A |
| Lin5-Cer3.2 | Deletion of a T and insertion of a GCG sequence | Deletion of an A |
| Lin5-Cer3.3 | Deletion of a T and insertion of a GCG sequence | Deletion of an A |
| Lin5-Cer3.4 | Deletion of a T and insertion of a GCG sequence | Deletion of an A |
| Lin5-Cer3.5 | Deletion of a T and insertion of a GCG sequence | Deletion of an A |
| Lin5-Cer3.6 | Deletion of a T and insertion of a GCG sequence | Deletion of an A |
| Lin5-Cer3.7 | Deletion of a T and insertion of a GCG sequence | Deletion of an A |
| Lin5-Cer3.8 | Deletion of a T and insertion of a GCG sequence | Deletion of an A |
| Lin5-Cer3.9 | Deletion of a T and insertion of a GCG sequence | Deletion of an A |
| Lin5-Cer3.10 | Deletion of a T and insertion of a GCG sequence | Deletion of an A |
| Lin5-Cer3.11 | Deletion of a T and insertion of a GCG sequence | Deletion of an A |
| Lin5-Cer3.12 | Deletion of a T and insertion of a GCG sequence | Deletion of an A |
| Lin5-Cer3.13 | Deletion of a T and insertion of a GCG sequence | Deletion of an A |
| Lin5-Cer3.14 | Deletion of a T and insertion of a GCG sequence | Deletion of an A |
| Lin5-Cer3.15 | Deletion of a T and insertion of a GCG sequence | Deletion of an A |
| Lin5-Cer3.16 | Deletion of a T and insertion of a GCG sequence | Deletion of an A |
| Lin5-Cer3.17 | Deletion of a T and insertion of a GCG sequence | Deletion of an A |
| Lin5-Cer3.18 | Deletion of a T and insertion of a GCG sequence | Deletion of an A |
| Lin5-Cer3.19 | Deletion of a T and insertion of a GCG sequence | Deletion of an A |
| Lin5-Cer3.20 | Deletion of a T and insertion of a GCG sequence | Deletion of an A |
| Lin5-Cer3.21 | Deletion of a T and insertion of a GCG sequence | Deletion of an A |
| Lin5-Cer3.22 | Deletion of a T and insertion of a GCG sequence | Deletion of an A |
| Lin5-Cer3.23 | Deletion of a T and insertion of a GCG sequence | Deletion of an A |
| Lin5-Cer3.24 | Deletion of a T and insertion of a GCG sequence | Deletion of an A |
| Lin5-Cerv4.1 | Scrambled sequence | Scrambled sequence |
| Lin5-Cerv4.2 | Scrambled sequence | Scrambled sequence |
| Lin5-Cerv4.3 | Deletion of a T | - |
| Lin5-Cerv4.4 | Deletion of a T | Deletion of a T |
| Lin5-Cerv4.5 | Scrambled sequence | Scrambled sequence |
| Lin5-Cerv4.6 | Deletion of a T | Deletion of a T |
| Lin5-Cerv4.7 | Scrambled sequence | Scrambled sequence |
| Lin5-Cerv4.8 | Scrambled sequence | Scrambled sequence |
| Lin5-Cerv4.9 | Deletion of a T | Deletion of a T |
| Lin5-Cerv4.10 | Scrambled sequence | Scrambled sequence |
| Lin5-Cerv4.11 | Deletion of a T | Deletion of a T |
| Lin5-Cerv4.12 | Deletion of a T | Deletion of a T |
| Lin5-Cerv4.13 | Deletion of a T | - |
| Lin5-Cerv4.14 | Scrambled sequence | Scrambled sequence |
| Lin5-Cerv4.15 | Scrambled sequence | Scrambled sequence |
| Lin5-Cerv4.16 | Scrambled sequence | Scrambled sequence |
| Lin5-Cerv4.17 | Deletion of a T | Deletion of a T |
| Lin5-Cerv4.18 | Deletion of a T | - |
| Lin5-Cerv4.19 | Deletion of a T | Deletion of a T |
| Lin5-Cerv4.20 | Scrambled sequence | Scrambled sequence |
| Lin5-Cerv4.21 | Scrambled sequence | Scrambled sequence |
| Lin5-Cerv4.22 | Deletion of a T | Deletion of a T |
| Lin5-Cerv4.23 | Deletion of a T | Deletion of a T |
| Lin5-Cerv4.24 | Deletion of a T | Deletion of a T |
| T Cervil Lin5 |  |  |
| Lin5-Cer2.1 | Deletion of a T and insertion of a GCG sequence | Deletion of an A |
| Lin5-Cer2.2 | Deletion of a T and insertion of a GCG sequence | Deletion of an A |
| Lin5-Cer2.3 | Deletion of a T and insertion of a GCG sequence | Deletion of an A |
| Lin5-Cer2.4 | Deletion of a T and insertion of a GCG sequence | Deletion of an A |
| Lin5-Cer2.5 | Deletion of a T and insertion of a GCG sequence | Deletion of an A |
| Lin5-Cer2.6 | Deletion of a T and insertion of a GCG sequence | Deletion of an A |
| Lin5-Cer2.7 | Deletion of a T and insertion of a GCG sequence | Deletion of an A |
| Lin5-Cer2.8 | Deletion of a T and insertion of a GCG sequence | Deletion of an A |
| Lin5-Cer2.10 | Deletion of a T and insertion of a GCG sequence | Deletion of an A |
| Lin5-Cer2.11 | - | - |
| Lin5-Cer2.12 | Deletion of a T and insertion of a GCG sequence | Deletion of an A |
| Lin5-Cer2.13 | Deletion of a T and insertion of a GCG sequence | Deletion of an A |
| Lin5-Cer2.14 | Deletion of a T and insertion of a GCG sequence | Deletion of an A |
| Lin5-Cer2.15 | Deletion of a T and insertion of a GCG sequence | Deletion of an A |
| Lin5-Cer2.16 | Deletion of a T and insertion of a GCG sequence | Deletion of an A |
| Lin5-Cer2.17 | Deletion of a T and insertion of a GCG sequence | Deletion of an A |
| Lin5-Cer2.18 | Deletion of a T and insertion of a GCG sequence | Deletion of an A |
| Lin5-Cer2.19 | Deletion of a T and insertion of a GCG sequence | Deletion of an A |
| Lin5-Cer2.20 | - | - |
| Lin5-Cer2.21 | Deletion of a T and insertion of a GCG sequence | Deletion of an A |
| Lin5-Cer2.22 | Deletion of a T and insertion of a GCG sequence | Deletion of an A |
| Lin5-Cer2.23 | Deletion of a T and insertion of a GCG sequence | Deletion of an A |
| Lin5-Cer2.24 | Deletion of a T and insertion of a GCG sequence | Deletion of an A |
| Lin5-Cer2.25 | Deletion of a T and insertion of a GCG sequence | Deletion of an A |
| Lin5-Cer2.26 | Deletion of a T and insertion of a GCG sequence | Deletion of an A |
| Lin5-Cer2.27 | Deletion of a T and insertion of a GCG sequence | Deletion of an A |

| **Name** | **Mutation on gRNA1** | **Mutation on gRNA2** |
| --- | --- | --- |
| T Levovil Suc3 |  |  |
| Suc3 Lev1.1 | Deletion of a CCTTGACACCAA sequence | Deletion of an A |
| Suc3 Lev1.2 | Deletion of a CCTTGACACCAA sequence | Deletion of an A |
| Suc3 Lev1.3 | Deletion of a CCTTGACACCAA sequence | Deletion of an A |
| Suc3 Lev1.4 | Deletion of a CCTTGACACCAA sequence | Deletion of an A |
| Suc3 Lev1.5 | Deletion of a CCTTGACACCAA sequence | Deletion of an A |
| Suc3 Lev1.6 | Deletion of a CCTTGACACCAA sequence | Deletion of an A |
| Suc3 Lev1.7 | Deletion of a CCTTGACACCAA sequence | Deletion of an A |
| Suc3-Lev2.1 | Deletion of a CCTTGACACCAA sequence | Deletion of an A |
| Suc3-Lev2.2 | Deletion of a CCTTGACACCAA sequence | Deletion of an A |
| Suc3-Lev2.3 | Deletion of an AAGTCCTTGACACCAA sequence | Deletion of an AGT sequence |
| Suc3-Lev2.4 | Deletion of an AAGTCCTTGACACCAA sequence | Deletion of an AGT sequence |
| Suc3-Lev2.5 | Deletion of an AAGTCCTTGACACCAA sequence | Deletion of an AGT sequence |
| Suc3-Lev2.6 | Deletion of an AAGTCCTTGACACCAA sequence | Deletion of an AGT sequence |

| **Name** | **Mutation on gRNA1** | **Mutation on gRNA2** |
| --- | --- | --- |
| T Levovil FRK3 |  |  |
| FRK3-Lev2.1 | Deletion of a GA | - |
| FRK3-Lev2.2 | - | - |
| FRK3-Lev2.3 | Deletion of a GA | - |
| FRK3-Lev2.4 | Deletion of a GA | - |
| FRK3-Lev2.5 | Scrambled sequence | Scrambled sequence |
| FRK3-Lev2.6 | Deletion of a GA | - |
| FRK3-Lev2.7 | Scrambled sequence | Scrambled sequence |
| FRK3-Lev2.8 | Deletion of a GA | - |
| FRK3-Lev2.9 | Deletion of a GA | - |
| FRK3-Lev2.10 | Deletion of a GA | - |
| FRK3-Lev2.11 | Deletion of a GA | - |
| FRK3-Lev2.12 | - | - |
| FRK3-Lev3.1 | Substitution of T by C | - |
| FRK3-Lev3.2 | - | - |
| FRK3-Lev3.3 | Substitution of T by C | - |
| FRK3-Lev3.4 | Substitution of T by C | - |
| FRK3-Lev3.5 | heterozygous KO from 305bp | - |
| FRK3-Lev3.6 | heterozygous KO from 305bp | - |
| FRK3-Lev3.7 | - | - |
| FRK3-Lev3.8 | - | - |
| FRK3-Lev3.9 | - | - |

| T Levovil Lin5 |  |  |
| --- | --- | --- |
| Lin5 Lev1.1 | Deletion of a T | Deletion of a T |
| Lin5 Lev1.2 | Deletion of a T | Deletion of a T |
| Lin5 Lev1.3 | Deletion of a T | Deletion of a T |
| Lin5 Lev1.4 | Deletion of a T | Deletion of a T |
| Lin5 Lev1.5 | Deletion of a T | Deletion of a T |
| Lin5 Lev1.6 | Deletion of a T | Deletion of a T |
| Lin5 Lev1.7 | Deletion of a T | Deletion of a T |
| Lin5 Lev1.8 | Deletion of a T | Deletion of a T |
| Lin5 Lev1.9 | unreadable sequence | unreadable sequence |
| Lin5 Lev1.10 | unreadable sequence | unreadable sequence |
| Lin5-Lev2.1 | - | - |
| Lin5-Lev2.2 | - | - |
| Lin5-Lev2.3 | - | - |
| Lin5-Lev2.4 | Deletion of a T | Deletion of an AT |
| Lin5-Lev2.5 | - | - |
| Lin5-Lev2.6 | Deletion of a T | Deletion of an AT |
| Lin5-Lev2.7 | - | - |
| Lin5-Lev2.8 | - | - |
| Lin5-Lev2.9 | Deletion of a T | Deletion of an AT |
| Lin5-Lev2.10 | Scrambled sequence | Deletion of an AT |
| Lin5-Lev2.11 | Scrambled sequence | Deletion of an AT |
| Lin5-Lev2.12 | - | - |
| Lin5-Lev2.13 | Deletion of a T | Deletion of an AT |
| Lin5-Lev2.14 | Deletion of a T | Deletion of an AT |

**Table 6 : Transgenesis experiment 2018 results (number of explants, cell clusters and plants at each step****)**

| target gene Malate 30 | Number of explants in culture | Cell clusters | Buds | Isolated plants | Acclimated plants | Mutant lines obtained |
| --- | --- | --- | --- | --- | --- | --- |
| Accession Cervil | | | | | | |
| TGF β1 | 32 | 19 | 21 | 22 | 9 | 4 |
| Control | 32 | 9 | 19 | 10 | 10 | 0 |
| Accession Levovil | | | | | | |
| TGF β1 | 32 | 26 | 20 | 64 | 43 | 8 |
| Control | 32 | 20 | 13 | 16 | 4 | 4 |
| Accession Ferum | | | | | | |
| TGF β1 | 32 | 20 | 19 | 16 | 9 | 6 |
| Control | 32 | 12 | 15 | 10 | 4 | 3 |

**Table 7 : Transgenesis experiment 2019 (number of explants, cell clusters and plants at each step)**

| target gene Malate 30 | Number of explants in culture | Cell clusters | Buds | Isolated plants | Acclimated plants | Mutant lines obtained |
| --- | --- | --- | --- | --- | --- | --- |
| Accession Cervil | | | | | | |
| TGF β1 | 48 | 71 | 59 | 67 | 33 | 16 |
| Control | 48 | 41 | 15 | 20 | 0 | 0 |
| Accession Levovil | | | | | | |
| TGF β1 | 48 | 49 | 42 | 46 | 27 | 18 |
| Control | 48 | 36 | 20 | 9 | 12 | 5 |
| Accession Stupike | | | | | | |
| TGF β1 | 48 | 33 | 49 | 59 | 42 | 20 |
| Control | 48 | 37 | 9 | 10 | 21 | 9 |
| Accession Criollo | | | | | | |
| TGF β1 | 48 | 33 | 69 | 53 | 60 | 49 |
| Control | 48 | 41 | 32 | 37 | 24 | 18 |
| Accession LA1420 | | | | | | |
| TGF β1 | 48 | 50 | 51 | 64 | 84 | 44 |
| Control | 48 | 37 | 25 | 36 | 63 | 30 |

**Table 8 : Transgenesis experiment 2022 (number of explants, cell clusters and plants at each step)**

| Malate 30 Knock-in target gene for Cervil accession | Number of explants in culture | Cell clusters | Buds | Isolated plants | Acclimated plants | Mutant lines obtained |
| --- | --- | --- | --- | --- | --- | --- |
| Experiment 1 | | | | | | |
| TGF β1 | 60 | 55 | 75 | 19 | 11 | 0 polluted box |
| IL-1β | 60 | 60 | 69 | 15 | 8 | 5 |
| EGF | 60 | 64 | 41 | 10 | 3 | 1 |
| Control | 60 | 25 | 25 | 1 | 0 | 0 |
| Experiment 2 | | | | | | |
| TGF β1 | 60 | 96 | 59 | 36 | 27 | 28 |
| IL-1β | 60 | 85 | 82 | 37 | 36 | 33 |
| EGF | 60 | 84 | 57 | 27 | 8 | 5 |
| Control | 60 | 21 | 23 | 5 | 2 | 0 |
| Experiment 3 | | | | | | |
| TGF β1 | 60 | 79 | 51 | 26 | 28 | 33 |
| IL-1β | 60 | 108 | 72 | 37 | 40 | 25 |
| EGF | 60 | 78 | 30 | 17 | 9 | 9 |
| Control | 60 | 27 | 18 | 4 | 3 | 0 |
